# Computational rheometry of viscoelastic networks: From random graphs to biomolecular condensates

**DOI:** 10.1101/2025.08.20.671282

**Authors:** Ruoyao Zhang, Gaurav Mitra, Souradeep Ghosh, Rohit V. Pappu

## Abstract

Multivalent biomacromolecules including multi-domain and intrinsically disordered proteins form biomolecular condensates via reversible phase transitions. Condensates are viscoelastic materials that display composition-specific rheological properties and responses to mechanical forces. Graph-based descriptions of microstructures can be combined with computational rheometry to model the outcomes of passive and active mechanical measurements. We consider two types of network models for microstructures. In the Jeffreys model, each edge in the network is a Jeffreys element. In the Stokes-Maxwell model, each edge is a Maxwell element that is embedded in an incompressible viscous fluid that can undergo Stokes flow. We describe results from comparative assessments of the two models for individual elements, ordered lattices, random geometric graphs, structured graphs, and graphs for condensates that are extracted from coarse-grained simulations of disordered proteins. Results from deformation and relaxation tests and flow field analysis reveal how distinct length and time scales contribute to the responses of different types of networks. No single test provides definitive assessments of the connections between material properties and microstructures. Instead, a range of active and passive rheometric tests are essential for distinguishing the responses of different types of networks. Our work establishes computational rheometry as a framework for bridging disparate length and timescales to assess how molecular-scale interactions and dynamics give rise to viscoelastic responses on the mesoscale.

## I. INTRODUCTION

Condensation of multivalent biomacromolecules drives the formation of compositionally distinct membraneless bodies known as biomolecular condensates [1–4]. The molecular drivers of condensation span diverse classes. They include multidomain proteins with modular interaction sites [2, 5, 6], intrinsically disordered proteins (IDPs) whose phase behavior emerges from specific sequence grammars [7–13], and multivalent nucleic acids [14–16]. The interplay between molecular-scale thermal fluctuations, multivalent interactions, and solvent reorganization gives rise to emergent mesoscale properties that distinguish condensates from homogeneous one-phase solutions [6, 13, 17–20]. In addition to featuring spatial inhomogeneities across multiple length scales [6, 18, 21–25], condensates exhibit complex, time-dependent mechanical responses [26–29].

Many condensates display viscoelastic behaviors mirroring the behaviors of many soft matter systems [30]. They can respond as elastic solids over short timescales while flowing as viscous liquids over longer periods [23, 27, 31–41]. This allows condensates to generate and respond to mechanical forces through bulk elasticity and interfacial free energies [27, 29, 33, 42–44]. The interplay between dominantly viscous versus elastic behaviors can change with time [26, 28] or applied shear [45]. As an example of the latter, Shen et al.,[45] showed that condensates can undergo shear-mediated transitions from dominantly viscous to dominantly elastic behaviors, whereby elasticity dominates and long timescales even though viscosity dominates over short timescales.

Different experiments have been designed and deployed to probe the rheological and interfacial properties of condensates across different length and time scales [46, 47]. Fluorescence recovery after photobleaching provides comparative inferences regarding molecular transport within condensates [48]. Fluorescence correlation spectroscopy provides assessments of translational diffusion of individual molecules within condensates [10, 49–51]. Förster resonance energy transfer measurements quantify chain reconfiguration times within condensates [50, 51]. Information regarding intermediate length and time scales can be accessed via methods that quantify inverse capillary velocities [31, 52] and time-correlated motions of probes within condensates. For the latter, different techniques have been deployed and these include active microrheology with optical trapping [26, 28], passive microrheology with and without optical trapping [28, 36, 37], dynamic light scattering [41], and atomic force microscopy [29, 53]. On the scale of condensates as a whole, micropipette aspiration provides inferences regarding interfacial tension and yield stress [54]. Finally, active microrheology helps assess the responses of condensates to applied forces using creep tests, measuring speeds of fusion, and rearrangement dynamics of condensates [28, 34, 35].

Condensation combines reversible binding [55], reversible networking transitions known as physical gelation or bond percolation [2, 56–60], and density transitions or demixing of incompatible molecules that drive phase separation [1, 2, 61]. The synergies among these processes can be captured in coarse-grained simulations that provide insights regarding the molecular scale interactions, structures, and dynamics that contribute to the emergence of inhomogeneities and two-phase behavior on the mesoscale [57, 58, 62–78]. All atom simulations with explicit representations of solvent molecules also hold significant promise, although their reach is limited in terms of length and timescales that can be accessed [50, 79–81]. Continuum models allow for phenomenological descriptions of viscoelasticity and the effects of hydrodynamic interactions on specific length and time scales [82], but these approaches lack information regarding the structural details on the molecular and mesoscales.

Discerning how condensates function as mechanosensitive and mechano-responsive materials requires a complete characterization of viscoelasticity and mechanical responses across a broad spectrum of length and time scales that are relevant to the functions of condensates [19, 27, 42, 44, 78]. What is needed is a framework that can be complemented with coarse-grained or atomistic simulations, whereby the structures of condensate interiors, extracted from simulations, are represented as graphs [83], so that passive and active time-dependent and time-independent responses of these graphs can be computed. The rheometric tests of interest include: (i) time-dependent in-phase (elastic) and out-of-phase (viscous) responses to oscillatory shear stress; and (ii) time-dependent responses to constant shear stress, constant strain, and constant strain rates. Here, we present our adaptation and generalization of a formalism known as computational rheometry that was introduced by Wróbel et al., [84] for modeling viscoelastic networks.

A key consideration in computational rheometry is capturing hydrodynamic interactions. At the mesoscale, flows typically occur at low Reynolds numbers [85], where inertial effects are negligible and viscous forces dominate.

For condensates of characteristic size *L* ≲ 1 µm, velocity *v* ~ 0.01-1 µm/s, and solvent viscosity *µ* comparable to water [86, 87], the Reynolds number Re = *ρvL/µ* ≪ 1 implies a regime governed by Stokes drag and long-range hydrodynamic coupling, which can mediate collective motion across the condensate. The significance of hydrodynamic effects is further characterized by the Péclet number Pe = *vL/D*, where *D* is the effective diffusion coefficient of the condensate as a whole [86, 88]. When Pe ≳ 1, advective transport rivals diffusion, producing non-equilibrium phenomena such as shape deformation, shear alignment, and internal circulation. The viscoelastic timescale of a material, *τ*_v_ ~ *η/G* (with *η* and *G* denoting the viscous and shear modulus), defines the crossover between predominantly elastic (*t* ≪ *τ*_v_) and viscous (*t* ≫ *τ*_v_) responses. Multiple such timescales may exist in complex fluids, each associated with distinct relaxation mechanisms. These timescales can overlap or compete with hydrodynamic relaxation times, generating a rich interplay between internal viscoelasticity and solvent-mediated dissipation [89].

In what follows, we compare the responses of networks of Jeffreys elements in a vacuum to networks of Maxwell elements under Stokes flow. The latter is referred to as a network of Stokes-Maxwell elements (Fig. 1). We model the responses of random geometric graphs (RGGs) and compare these to responses of ordered lattices. This shows that RGGs are equivalent to ordered lattices, with the degree of connectivity being the only determinant of mechanical responses in both cases. However, the mechanical responses of networks become sensitive to spatial organization within networks when we consider structured graphs. These findings motivate the extraction of spatially embedded graphs from coarse-grained simulations of condensates, which we use to model viscoelasticity and the coupling of networks to Stokes flow.

**FIG. 1.**
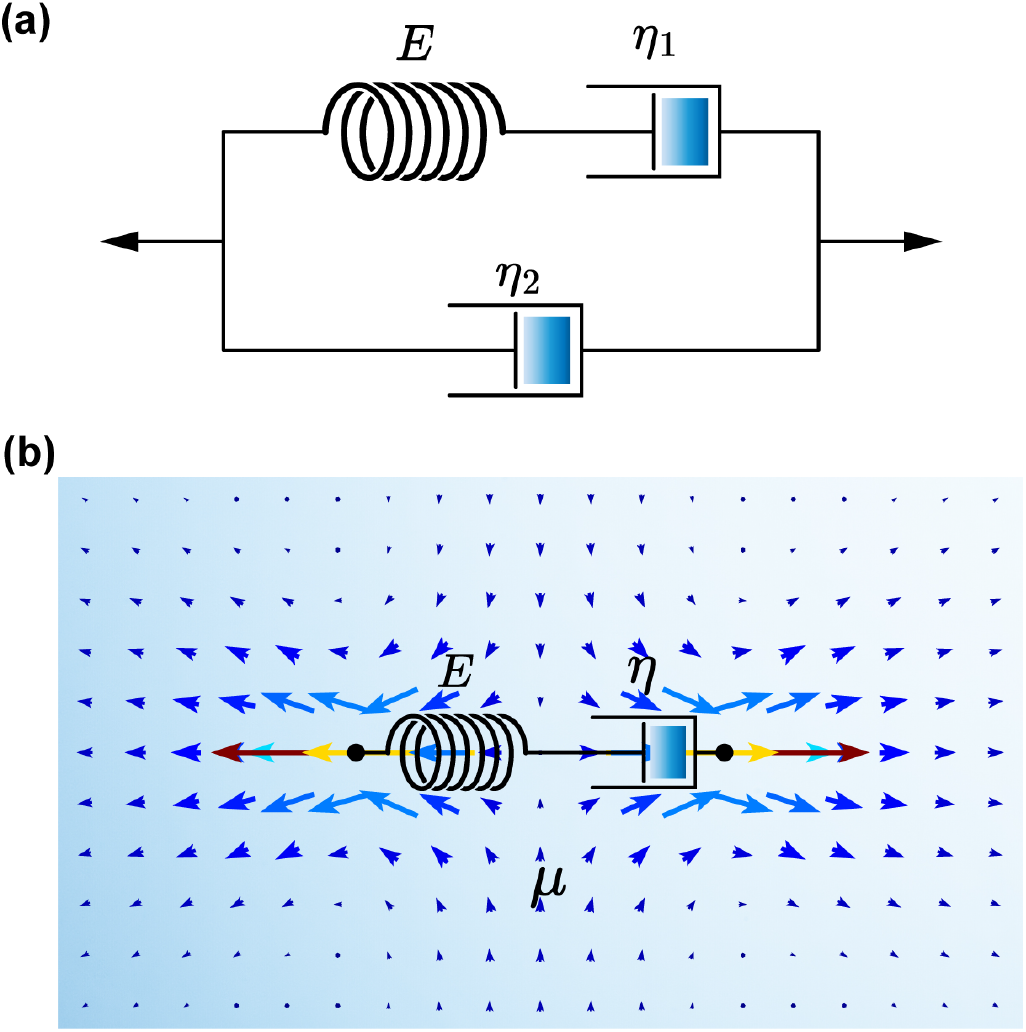
Schematics of (a) a single Jeffreys element and (b) a single Stokes-Maxwell element. A Jeffreys element comprises a spring of elastic modulus *E* that is in series with a dashpot of viscosity *η*_1_. It also includes a second dashpot in parallel that has a viscosity *η*_2_. The Stokes-Maxwell element comprises a Maxwell element of elastic modulus *E* and intrinsic viscosity *η* that is coupled via Stokes flow to a fluid of viscosity *µ*. Blue arrows indicate flow directions when the element is subjected to tensile force.

## II. MODELS FOR VISCOELASTIC NETWORKS

At the outset, we assess the contributions due to the surrounding solvent and develop quantitative criteria that allow us to adjudicate when solvent effects must be accounted for to model the rheological responses of a viscoelastic network. We follow the linearized two-fluid analysis of Levine and Lubensky [90, 91], in which a viscoelastic network is viscously coupled via a friction coefficient Γ to an incompressible Newtonian fluid. We derive a hydrodynamic screening length *ξ*(*ω*) from the reduced two-fluid model for a viscoelastic network in a Stokes fluid (detailed in Appendix A). We obtain the relation,

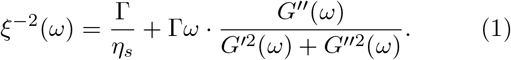

Here, *η*_*s*_ is the solvent viscosity, *ω* is the frequency corresponding to the inverse of the timescale at which the measurement is made, and *G*′ and *G*″ are the real (in-phase) and imaginary (out-of-phase) parts of the complex shear modulus. We define a dimensionless coupling parameter ℋ (*ω*) = *b/ξ*(*ω*), where *b* is the characteristic mesh size of the network. The strong hydrodynamic screening limit corresponds toℋ(*ω*) ≫ 1. In this limit, hydrodynamic interactions are strongly screened such that the flows decay within one mesh size. If we consider each edge of the network to be a Maxwell element, it is equivalent to adding a parallel dashpot to each edge, as each edge feels only its local drag. Therefore, the entire network can be modeled as a Jeffreys network without explicit considerations of the fluid.

In the strong hydrodynamic coupling limit, ℋ (*ω*) ≪ 1, the screening length exceeds many mesh cells. A force on one node generates long-range solvent motion that couples the entire network. In this scenario, the network can be viewed as being immersed in a fluid under Stokes flow, and modeling each edge as a Maxwell element becomes more physically meaningful [92, 93] (Fig. 1b).

For biomolecular condensates, within frequency ranges that are accessible to experiments [41], we estimate that 1 *<* ℋ *<* 2 (Appendix A). This suggests that condensates lie between the strongly screened and strong hydrodynamic coupling limits. Therefore, we cannot ignore explicit fluid-network coupling in the context of modeling or measuring the viscoelastic properties of condensates. We propose that quantitative assessments of mechanical responses of condensates must use models pertaining to the two limits *viz*., the Jeffreys and Stokes-Maxwell models, and probe system-specific parameter sensitivities imposed by these models. This requires a framework to either analyze experimental data by simulating the responses and parameter sensitivities of different models, identifying the regimes that best describe the data, or bridging all-atom or coarse-grained simulations of condensates to the length and timescales probed in experiments. Here, we construct, analyze, and compare dynamic moduli and mechanical responses for different types of graphs modeled as edge-based Jeffreys networks or hybrid node-and-edge-based Stokes-Maxwell networks. Both models share the same spatial network topology defined by undirected edges ℰ between nodes/vertices 𝒱; only the treatment of hydrodynamic effects differs. Our description and usage of both models rely on non-dimensionalized parameters. Appendix B provides a prescription for converting between dimensional and non-dimensional parameters.

### A. Edge-based Jeffreys networks

Viscoelastic materials exhibit a hybrid mechanical response between purely elastic (Hookean) and purely viscous (Newtonian) behaviors. One way to study viscoelasticity numerically is to discretize the material into a network of nodes and edges, with a constitutive law for each node / edge. A Jeffreys element, consisting of a Maxwell branch in parallel with an additional dashpot (Fig. 1a), provides a useful starting point. It captures the essence of viscoelasticity while also being derivable from microscopic considerations [94].

We consider a spatial network with each edge being a Jeffreys element that follows the constitutive relation:

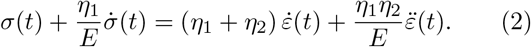

Here, *σ*(*t*) is the stress, *ε*(*t*) is the strain, *E* is the elastic modulus, and *η*_1_, *η*_2_ are the viscosities of the dashpots in the Maxwell and parallel branches, respectively. Switching to the frequency domain with a factor *e*^*iωt*^ yields a stress-strain relationship,

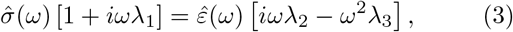

where *λ*_1_ = *η*_1_*/E, λ*_2_ = *η*_1_ + *η*_2_ and *λ*_3_ = *η*_1_*η*_2_*/E*. This leads to the following expression for the complex shear modulus,

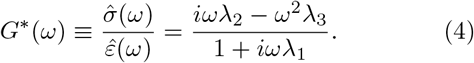

When each Jeffreys element is subjected to small amplitude oscillatory shear on the boundary nodes, we can solve for *G*^*^(*ω*) = *G*′ + *iG*″, where *G*′ = *Eη*^2^*ω*^2^*/*(*E*^2^ +*η*^2^*ω*^2^) and 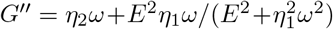 are the storage and loss moduli, respectively.

As detailed in Appendix C, we develop a frequency-domain analysis to calculate the dynamic moduli of undirected networks with each edge being a Jeffreys element. This formulation leads to the result that the shapes of the frequency dependencies of the dynamic moduli *G*′ and *G*″ are universal for every 3D Jeffreys network. The two curves translate vertically, and in concert, when changing the node positions or edge density.

### B. Network of Maxwell elements in a Stokes fluid

The Rouse model for dynamics of a single polymer in a dilute solution laid the groundwork for viscoelasticity of polymer solutions [92]. In this model, which is directly relevant for condensates formed by IDPs [51, 83], an *N* - bead polymer features *N* relaxation modes, and each mode is a Maxwell element comprising a spring and dash-pot in series. Thus, a polymer solution may be viewed as a generalized Maxwell system with Maxwell elements assembled in parallel [92]. While the Rouse model assumes a freely draining limit, the Zimm model accounts for the contributions of hydrodynamic interactions to the relaxation of each of the Maxwell elements [93]. Generalizing the Rouse-Zimm picture, one can view a viscoelastic material as a spatially organized network of Maxwell elements coupled to a fluid undergoing Stokes flow. Analyzing the mechanical responses of such a system requires a numerical approach and this is the computational rheometry formalism of of Wróbel et al., [84]. Using this formalism, we assume that a network of Maxwell elements is suspended in a Newtonian, purely viscous solvent. We then solve for the incompressible Stokes equations:

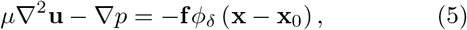

and

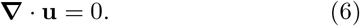

Here, **u**, *p*, and **f** are velocity, pressure, and force, respectively. *ϕ*_*δ*_ is a regularizing function (Eq. D1), which is a smooth approximation of a delta function. We need *δ* ≪ ∥**x**_*j*_ − **x**_*i*_∥ so that neighboring nodes do not overlap; otherwise the solvent can flow through the edges and the network becomes under-damped. The exact solution can be formulated as [84]

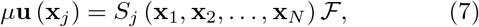

where *µ* is the fluid viscosity, ℱ is the force vector, and *S*_*j*_ is a regularized Stokeslet. The method of a regularized Stokeslet provides a way to circumvent the singular solution of the Stokes equation with a point force [95].

Immersing a Maxwell element in a fluid (Fig. 1b) leads to a distinct type of element that we refer to as a Stokes-Maxwell element [84]. The stress-stain relation of a Stokes-Maxwell element with time-dependent length 𝓁 follows

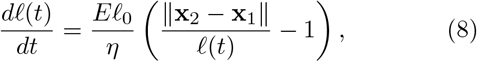

where **x**_1_(*t*) and **x**_2_(*t*) are the two endpoints of the element. We follow the protocols of Wróbel et al. [84], and details of the numerical methods are discussed in Appendix D.

## III. RESULTS

This section is organized progressively, advancing from simple, ordered networks to complex, random networks. Throughout, we use dimensionless parameters for *E, η*_1_, and *η*_2_ for Jeffreys elements and *E, η*, and *µ* of Stokes-Maxwell elements. All of our results are presented in terms of dimensionless moduli. To enable direct comparisons between the mechanical responses of edge-based Jeffreys networks and the nodal Stokes-Maxwell networks, we set *η*_1_ = *η* and *η*_2_ = *µ*.

### A. A single element

We first compare the viscoelastic responses of a single Jeffreys element in a vacuum with those of a single Stokes-Maxwell element. dynamic moduli were calculated analytically for the Jeffreys element (Eq. 4), and determined through small-amplitude oscillatory shear tests for the Stokes-Maxwell element. In Fig. 2a, we establish a baseline by setting all dimensionless parameters *E, η*, and *µ* equal to 1. The two methods show excellent agreement throughout the frequency range, with no crossover behavior. A single crossover emerges when *E* = *η* = 10^5^ while *µ* = 1 (Fig. 2b). Under these conditions, viscous forces from the Stokes fluid or the viscosity of the parallel dashpot in the Jeffreys model become negligible compared to the dashpot resistance in the Maxwell element, yielding characteristic Maxwell-like behavior. When *E* = *µ* = 1 and *η* = 10^5^ (Fig. 2c), the dynamic moduli exhibit Kelvin-Voigt-like behavior [84], as the highly viscous dashpot in the Maxwell element dominates the response.

**FIG. 2.**
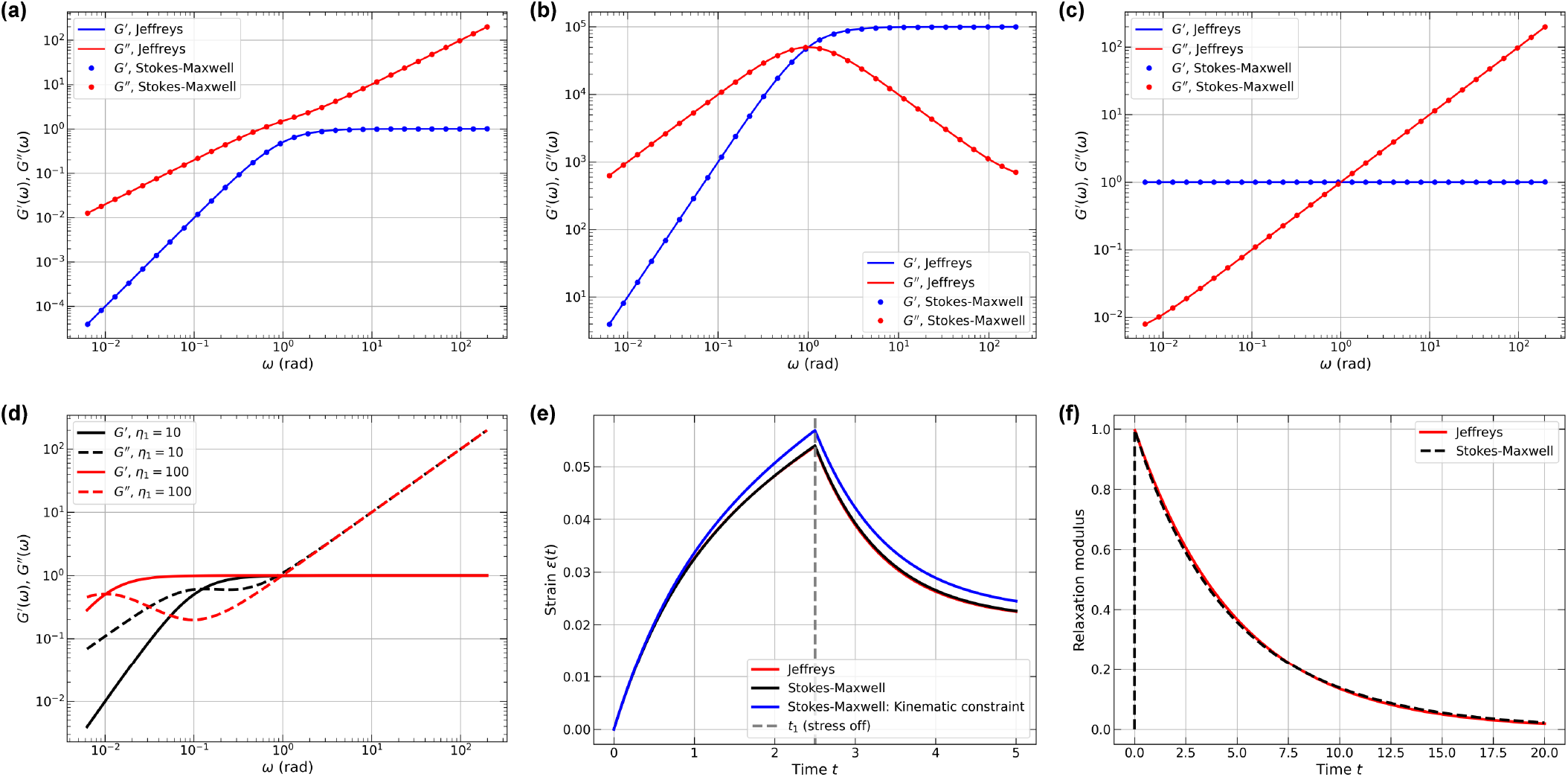
Viscoelastic responses of Jeffreys and Stokes-Maxwell elements probed in different limits. Dimensionless storage and loss moduli for (a) *E* = *η* = *µ* = 1, (b) *E* = *η* = 10^5^ and *µ* = 1, and (c) *E* = *µ* = 1 and *η* = 10^5^. Lines denote analytical solutions from the Jeffreys model and circles denote numerical solutions of a Stokes-Maxwell element. Blue: *G*′(*ω*); red: *G*″(*ω*). (d) Dimensionless storage and loss moduli when *η* increases from 10 (black) to 100 (red), with *E* = *µ* = 1. Solid and dashed lines represent *G*′(*ω*) and *G*″(*ω*), respectively. (e) Strain response during a creep test. The time *t*_1_ represents the time point when the force is removed. Red: analytical solution of the Jeffreys element; black: numerical solution of the Stokes–Maxwell element; blue: Stokes–Maxwell model with a kinematic constraint. Parameters: *E* = *µ* = 1, *η* = 5, and *f*_0_ = 0.05. (f) Relaxation modulus versus time. Red line: analytical solution of the Jeffreys element; black dashed line: numerical solution of the Stokes-Maxwell element. Parameters: *E* = *µ* = 1, *η* = 5, and *ε*_0_ = 0.1.

When *E* and *η* exceed *µ* by one to two orders of magnitude, single-element dynamic moduli exhibit a double-crossover behavior that is distinct from that of Maxwell or Kelvin-Voigt elements. By equating *G*′ and *G*″ and assuming *η* ≫ *µ*, we find that the high frequency crossover follows the relation *ω*_high_ = *E/µ*− *Eη*^−1^ − (*E* + *Eµ*) *η*^−2^+ *O*(*η*^−3^), and the low frequency crossover follows the relation *ω*_low_ = *Eη*^−1^ +2*Eµη*^−2^ +*O*(*η*^−3^). For fixed values of *E* and *µ*, these expressions reveal that *ω*_high_ remains rel-atively insensitive to *η*, while *ω*_low_ decreases significantly with increasing *η*. This prediction is confirmed in Fig. 2d, where double-crossover behavior emerges in a single element. As *η* increases from 10 to 100, the crossover region broadens considerably, with the lower crossover frequency shifting leftward by nearly an order of magnitude while the upper crossover remains relatively fixed. DLS measurements of coacervates formed by acidic peptides complexing with fluorescent proteins provide clear evidence of double crossover behavior and other frequency-dependent behaviors of *G*′ and *G*″ [41]. As shown here, the double crossover behavior can be explained using a single element. However, the observations of Fisher and Obermeyer [41] show additional complexities, requiring that we go beyond single elements. Fixing *η* = *µ* = 1 and increasing *E* from 10 to 100 leaves *G*″ essentially unchanged, whereas the plateau value for *G*′ shifts up, and the onset of the plateauing of *G*′ shifts to higher frequencies (data not shown).

For creep tests under step-stress loading, we derived analytical solutions for the Jeffreys element (see Appendix E) and obtained numerical solutions for the Stokes-Maxwell element. The test configuration fixes one node while applying a constant force to the other node that is parallel to the element axis. The response of the Stokes-Maxwell elements aligns perfectly with the analytical solution (Fig. 2e). We also examined an alternative boundary condition where the fixed node is constrained kinematically by prescribing zero velocity rather than solving for the required external forces. This approach produces a slight deviation from the analytical solution, highlighting the importance of consistent boundary condition implementation. Time-domain creep tests reveal how subtle differences in boundary conditions and hydrodynamic effects affect the mechanical responses. These differences are not apparent in the computed dynamic moduli or in the results of simple relaxation tests. The local strain-rate (or displacement) is not necessarily uniform in the Stokes-Maxwell element, nor is the internal stress distribution necessarily uniform in time. This breaks the uniform-strain assumption that enables analytical tractability in the Jeffreys element, demonstrating why numerical solutions become necessary when hydrodynamic interactions are included.

Fig. 2f compares the stress relaxation of single elements following a instantaneous strain. Note that this test does not involve moving point forces making it unlike the creep configuration. The analytical relaxation modulus for the Jeffreys element (see Appendix E) aligns closely with the Stokes-Maxwell result.

Having established the fundamental behavior of individual elements, we now examine more complex network structures, beginning with regular lattices. While single-element comparisons show close correspondence between the Jeffreys and Stokes-Maxwell formulations when we set *η*_2_ of the Jeffreys model to be equal to *µ*, the viscosity of the fluid, this correspondence breaks down in complex networks. When multiple nodes interact hydrodynamically, the Stokes-Maxwell formalism captures non-local dissipation effects that are absent in the Jeffreys model, thus leading to mechanical responses that are qualitatively different. Hence, for condensates, it becomes imperative to investigate both models and assess the parameter regimes that best describe the totality of the available data.

### B. Lattice networks

We consider a 5 × 5 × 5 cubic lattice network as shown in Fig. 3a. Each node within this network forms edges with its 18 nearest neighbors. We first compute the dynamic moduli of a Jeffreys network with this structure (Fig. 3b; see Appendix C). The nodes are partitioned into constrained and unconstrained portions based on the coordinates. Nodes in the bottom layer are fixed, the top layer is subjected to an oscillatory shear, and the interior nodes are unconstrained. Each element feels only its own spring and dashpots. So, when a larger lattice is sheared, every element contributes the same local stiffness/dissipation per row of elements. The result is that all edges share the same characteristic timescale and the same frequency-dependent impedance shape. The network stitches these impedances together but it does not introduce additional relaxation times. Changing the connectivity or the node degrees changes the net magnitude of the stiffness/damping, causing a vertical shift. Increasing the lengths of edges by a factor of *α* decreases the magnitudes of both dynamic moduli by a factor of 1*/α*^2^ (discussed in Appendix F).

**FIG. 3.**
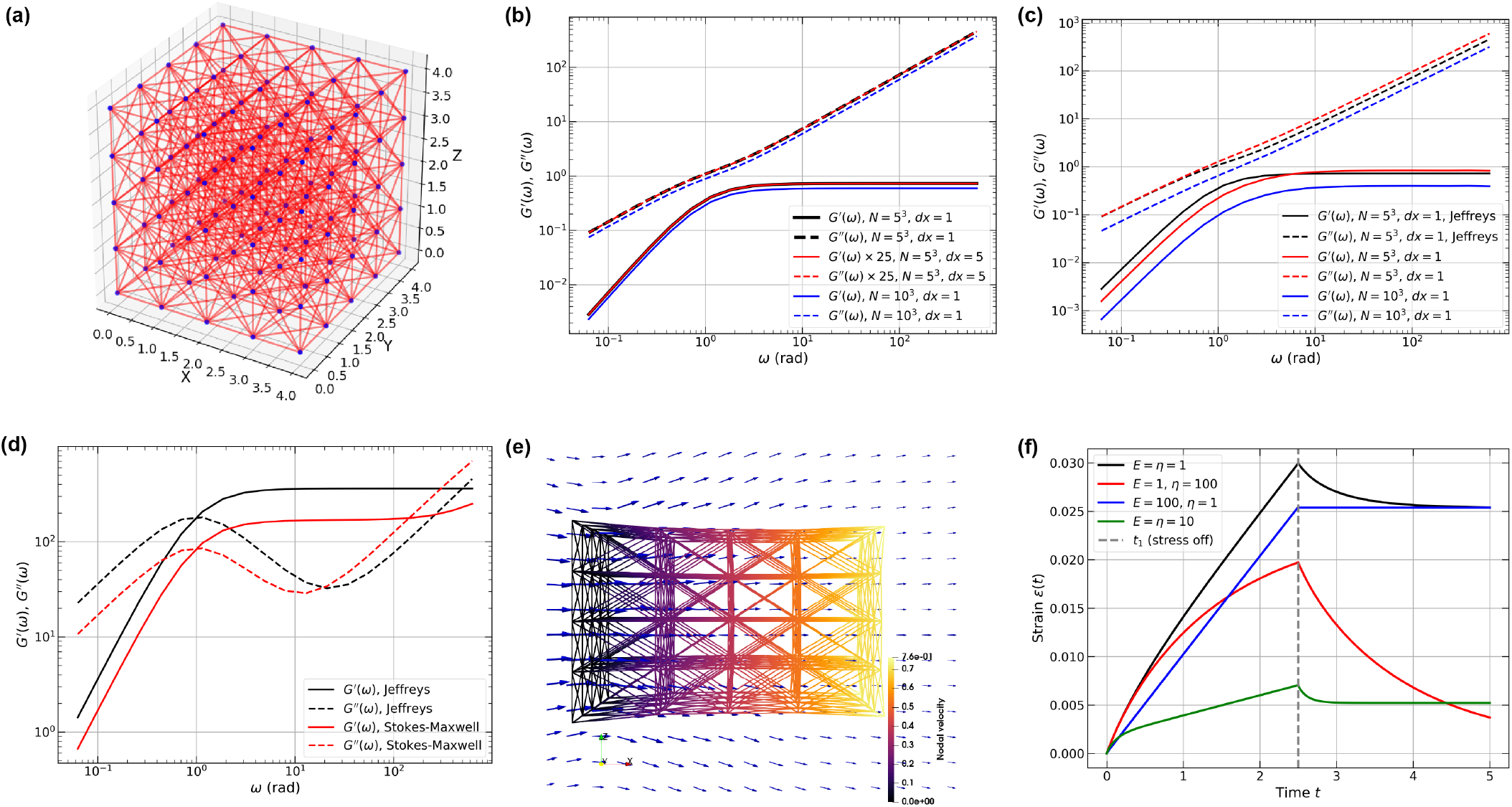
The viscoelastic responses of Jeffreys and Stokes-Maxwell elements at different limits. (a) Schematic of a 5 *×* 5*×* 5 lattice network with edges between nearest neighbors and diagonal nodes: *N* = 125 and *M* = 780. (b) Dimensionless dynamic moduli for a lattice network of Jeffreys elements with *E* = *η* = *µ* = 1. Black: 5 *×* 5 *×* 5 lattice with *dx* = 1; red: 5 *×* 5 *×* 5 lattice with *dx* = 5; blue: 10 *×* 10 *×* 10 lattice with *dx* = 1, here *N* = 1, 000 and *M* = 7, 560. (c) Dimensionless dynamic moduli of Stokes-Maxwell lattice networks with *E* = *η* = *µ* = 1. (d) dynamic moduli of the graph calculated using a Jeffreys network (*E* = *η* = 500, *µ* = 1) and a Stokes-Maxwell network (*E* = *η* = 500, *µ* = 1). (e) A representative snapshot of the Stokes-Maxwell lattice under large deformation during a creep test. Parameters: *E* = *η* = *µ* = 1, and *σ*_0_ = 1. Edges are colored by the dimensionless nodal velocity. Blue arrows denote fluid velocity in the cross-section area in the x-z plane. (f) Total strain response of the lattice network under creep tests with various physical conditions. Here, *t*_1_ represents the time for removing external driving forces. We set *σ*_0_ = 0.1.

When the size of the network increases to 10×10×10, the overall dynamic moduli become size-independent with a constant node spacing *dx* = 1. A slight deviation from the smaller lattice network is caused by the boundary effects. Further increases in size do not lead to any additional shifts of the curves.

The Stokes-Maxwell model on lattice networks behaves differently from a Jeffreys network. This is because every node couples to every other node through the Stokeslet kernel of the fluid. Fig. 3c shows that the Stokes-Maxwell network produces dynamic moduli that are different from the equivalent Jeffreys network. They share a similar plateau value for *G*′ at high frequencies, and equal *G*″ values at low frequencies. When we partition layers of nodes as in the Jeffreys case, the network gives a net top-plane reaction force

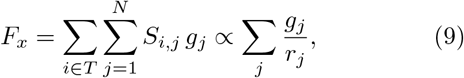

where *g*_*j*_ is the imposed velocity of node *j* and *r*_*j*_ its distance to the shear plane. Nodes deep inside the sample contribute weakly, which leads to a smaller pre-factor as the average over 1*/r*_*j*_ value drops. As a result, an amplified downshift of the dynamic moduli is observed in Fig. 3c when increasing the size of the lattice network to 10 ×10 ×10 while keeping the local edge density constant (*dx* = 1). The size-dependent hydrodynamic coupling in the Stokes-Maxwell network departs from the size-independent dynamic moduli in the corresponding Jeffreys network even without changing the network properties.

When we set *E* = *η* = 500, there is a clear downshift of both dynamic moduli (Fig. 3d). While the lower crossover frequencies are both at *ω* ≃ 1 rad, the higher cross-over frequency of the Stokes-Maxwell network exhibits a shift to lower values when compared to the Jeffreys network. Note that for the Stokes-Maxwell network, the value of *G*′ starts to deviate upward at high frequencies.

If each node is far from the walls and the network is sparse enough that inter-node hydrodynamic interactions are negligible, we may approximate the hydrodynamic damping as a single constant dashpot. In such an idealized scenario, the hydrodynamic drag is **F** = −6*πµδ***v**, and each node experiences a force that is decoupled from the velocity field, such that the Stokes-Maxwell network mimics a Jeffreys network in a vacuum. However, even such a system cannot produce a perfect match to the Jeffreys model at all frequencies. One can get close by increasing the size of the system, adding periodic or slip boundaries, or tuning parameters, but strict one-to-one equivalence is inhibited by the finite-size and boundary effects inherent to the discrete, localized fluid model.

Various active rheometric tests can be conducted on viscoelastic networks. These tests provide an important way to distinguish models that best describe the system of interest. We focus on the creep test using the Stokes-Maxwell model. As illustrated in Fig. 3e, we partition the nodes along the x-axis such that the left-most layer of nodes is fixed, while the right-most layer experiences a constant force on each node in the positive x-direction. The arrows indicate the flow field during the tensile motion of the network. The force is removed at dimensionless time *t*_1_ = 2.5 and one can compute the total strain evolution of the network as [84]

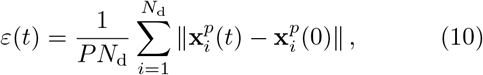

where *N*_d_ is the number of driven nodes, and *P* is the distance between two constrained layers. As the tensile forces are exerted, a fraction of the links gets stretched and the others are compressed. For the baseline case where *E* = *η* = *µ* = 1, the maximum total strain is largest and it is followed by a small recovery (Fig. 3f). Increasing *η* by two orders of magnitude leads to a lowered maximum total strain, and a strong relaxation towards the original configuration that mimics a Kelvin-Voigt-like creep response. In contrast, increasing *E* by two orders of magnitude produces a higher total strain. However, when the force is released, there is no recovery, and this is the response of a purely viscous system. Finally, when both *E* and *η* increase by an order of magnitude, the maximum total strain decreases further, accompanied by a rapid relaxation to a plateau value. Both the maximal strain achievable for a given force and the profile of relaxation upon releasing the force are relevant pieces of information that can be derived from a creep test. Thus, to use a creep test for drawing inferences regarding the relative magnitudes of *E* versus *η*, it becomes important to quantify the maximum strain and the relaxation profiles for different magnitudes of the applied force, while taking care to maintain the system in the linear response regime.

### C. Random geometric graphs in a spherical domain

The preceding discussion focused on the responses of ordered lattice networks [84]. This helped with comparisons between Jeffreys and Stokes-Maxwell networks. However, uniform lattices are poor approximations of condensates, which are exemplars of disordered mesoscale systems. As a step towards understanding networks of disordered systems, we investigated the responses of random graphs and applied the Jeffreys and Stokes-Maxwell formalisms to seek insights into their viscoelastic behaviors. These calibrations of random graphs are relevant in light of recent studies that have connected internal organization of condensates to network structures [70, 96–99], and the fact that condensates may be viewed as confined physical gels [25, 56, 57, 59–61]. We first investigated the responses of random geometric graphs (RGGs).

RGGs are graphs 𝒢 = (𝒱, ℰ) consisting of nodes 𝒱 placed in *d*-dimensional Euclidean space ℝ^*d*^, with edges ℰ (*X, Y*) ⊆ [𝒱]^2^ for *x* ∈ *X* and *y* ∈ *Y* added to connect pairs of points based on a cutoff distance ∥*Y*− *X*∥≤ *r*_*c*_ [100, 101]. The spatial positions of nodes are randomly generated within a spherical domain with a uniform volume distribution. The edges were constructed using the NetworkX python package [102]. For simplicity, we disallowed multiple edges between any two nodes and self-edges.

In Fig. 4a, for 200 random nodes within a spherical domain with *R* = 63.0, we show the network structure with a cutoff distance *r*_*c*_ = 30. The nodes with large degree centrality lie close to the center of the spherical domain. As *r*_*c*_ increases from 30 to 50, the total number of edges *M* increases from 1, 598 to 5, 880 (Fig. 4b), scaling as *M* ~ *r*^3^ (Fig. 4c; derived in Appendix G).

**FIG. 4.**
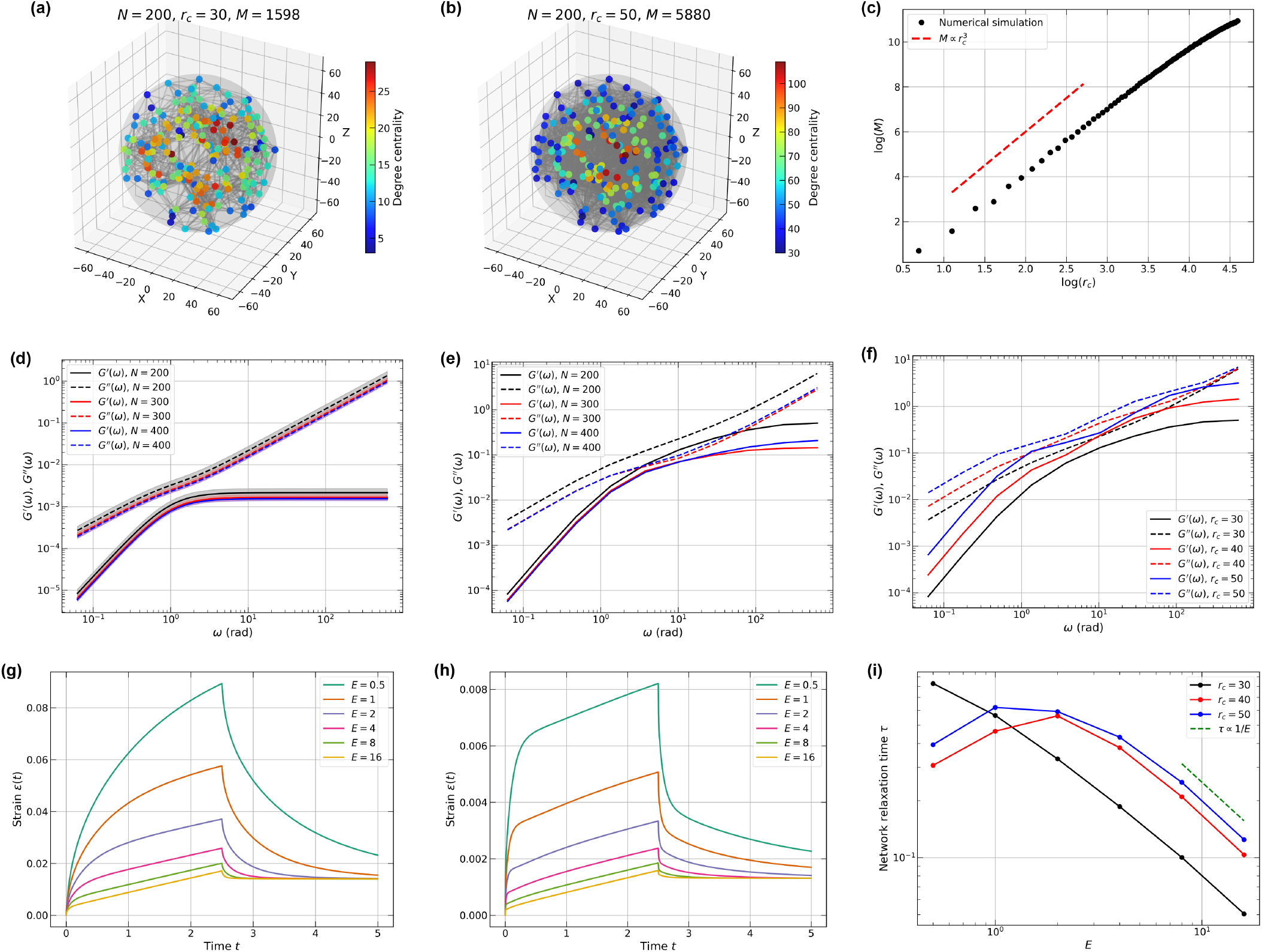
Viscoelastic responses of random geometric graphs (RGGs). (a,b) Representative RGGs with 200 nodes in a spherical domain of radius *R* = 63.0 at cutoff distances (a) *r*_*c*_ = 30 and (b) *r*_*c*_ = 50. (c) Number of edges as a function of cutoff distance *r*_*c*_ for *N* = 400 nodes in a spherical domain with *R* = 79.37. The red dashed line shows 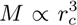 scaling. (d) Dimensionless dynamic moduli of Jeffreys networks with *N* = 200 (*R* = 63.0), 300 (*R* = 72.112), and 400 (*R* = 79.37) nodes. Parameters: *r*_*c*_ = 30, *E* = *η* = *µ* = 1. Shaded regions indicate standard errors. (e) dynamic moduli of the same networks as in (d) evaluated using the Stokes-Maxwell formalism. (f) Dimensionless dynamic moduli of a 200-node RGG at various cutoff distances using the Stokes-Maxwell model. (g,h) Creep tests for RGGs with (g) *r*_*c*_ = 30 and (h) *r*_*c*_ = 50. (i) Relaxation time *τ* as a function of cutoff distance. For all creep tests, *η* = 10 and *µ* = 1.

Unlike ordered lattice networks with flat faces and edges, random networks have irregular extrusions. We first establish the means to probe such networks by applying constant forces or strains to nodes in a confined volume. For example, by selecting the first and last 10% of the nodes in the x-direction, one can impose appropriate boundary conditions. Next, we keep the cutoff distance *r*_*c*_ and the average inter-node distances constant while varying the number of nodes to study the effect of network size. For a given node density in a spherical domain, the number of nodes *N* grows as *N* ∝ *R*^3^ with the domain radius *R*. We choose *N* = 100, *R* = 50.0 and *N* = 200, *R* ≈ 63.0 to maintain the average inter-node distance to be ~17.4. The dynamic moduli stay size-invariant for the Jeffreys networks with *N* = 200, 300, 400 as shown in Fig. 4d. Furthermore, we observe that varying the structures of the Jeffreys lattice network while assuming homogeneous bulk properties will only create vertical shifts in the dynamic moduli.

Introducing heterogeneous edge parameters will lead to multiple relaxation times throughout the network, which in turn results in different dynamic moduli. One can introduce ad hoc heterogeneities such as differences in edge weights. This can be used to generate numerical fits of the responses of RGGs to those that are measured. However, such an approach cannot shed light on whether a distinct network topology underlies the responses observed in a measurement. This requires tests based on active rheology such as the responses to creep tests, which are made possible using the Stokes-Maxwell model. Ac-cordingly, we evaluated RGGs using the Stokes-Maxwell model (Fig. 4e). A slight decrease in both dynamic moduli from *N* = 200 to 300 and 400 is observed due to the boundary effects. Larger networks tend to have converged dynamic moduli, which also agrees with the observations in lattice networks. As the value of *r*_*c*_ increases (Fig. 4f), the Stokes-Maxwell network with 200 nodes shows an increase in dynamic moduli as the number of edges increases, suggesting that the network becomes more elastic and more viscous at the same time for *ω <* 10^2^ rad. For higher frequencies, *G*″ values converge to the same curve and *G*′ plateaus at different values. We note that for Stokes-Maxwell RGGs, the value of *G*′ plateaus at higher frequencies than the corresponding Jeffreys network (cf. Fig. 4d) or the lattice networks.

To gain more insights into spherical RGGs, we performed creep tests with constant forces. We fixed the dimensionless viscosities *η* = 10 and *µ* = 1, and varied the value of *E* for the 200-node RGG. For low-connectivity, with *r*_*c*_ = 30, the maximum strain decreases as *E* increases (Fig. 4g). Thus, as the RGG becomes more elastic, the extent of tensile deformation becomes smaller because the total force exerted is constant. Additionally, the network with small values of *E* ranging from 0.5 to 2 can relax to more than half of the total strain for *t <* 5. It quickly plateaus after removal of the force for larger *E* values, indicating different relaxation times. Similarly, for the higher-connectivity case with *r*_*c*_ = 40 (Fig. 4h), the maximum strain of the network also decreases with increasing values of *E*. However, the magnitudes of the total strain become an order of magnitude smaller than the case with *r*_*c*_ = 30 (cf. Fig. 4g). Moreover, we note that the period of time (*t <* 2.5) that the network shows a constant strain rate becomes longer, which is a characteristic creep behavior of purely viscous materials.

Networks are characterized by a spectrum of relaxation times, and we estimate and compare the overall relaxation times of RGGs with different *r*_*c*_ values. An exponential decay function *ε*(*t*) = *a* + *b* exp(−*t/τ*) can be used to fit and extract the characteristic relaxation time *τ* of the networks for *t* ≥2.5. In Fig. 4i, which plots the relaxation times for networks with different *r*_*c*_ values, we observe that all cases conform to *τ* ∝1*/E* for large values of *E*. However, higher *r*_*c*_ cases exhibit a non-monotonic behavior at small values of *E*, deviating from what has been reported for lattice networks [103]. This non-monotonicity reflects contributions from stiff, cage-like structures that form for higher degrees of connectivity [104, 105].

Overall, despite being random in nodal positions and topology, the RGGs exhibit many similarities to lattice models. This is because the edges are constructed based purely on geometric cutoffs. To investigate edge constructions based on other criteria, we turned to random networks with spatial information.

### D. Spatially embedded Erdős-Rényi and small-world graphs

Many network models define edges without strictly enforcing a distance cutoff. We focus on two such networks: the Erdős-Rényi (ER) graph [106] and the small-world (SW) graph of Watts and Strogatz [107]. An identical set of *n* nodes is placed uniformly at random inside a spherical domain. Connectivity is assigned according to model-specific rules. For the spatially embedded ER networks 𝒢 (*n, p*), every unordered pair of nodes is connected independently with probability *p* ∈ [0, 1], mirroring the classical ER construction but entirely decoupled from Euclidean distance considerations. Multiple edges and self-loops are disallowed. In contrast, in spatially embedded SW networks, nodes are first labeled 0, …, *n*− 1 and connected in an index-based ring lattice, where each node is linked to the *k/*2 nodes immediately preceding and following it by label (not by Euclidean proximity). Each of these initial edges is then rewired with probability *β* ∈ [0, 1] following the Watts-Strogatz recipe, again without reference to the coordinates of the nodes. These approaches enable direct comparisons of distance-based connectivity (RGG), random connectivity (ER), and structured connectivity with random rewiring (SW).

Figs. 5a-c illustrate RGG, ER, and SW networks (*n* = 200) constructed from an identical set of node positions but with different connectivity rules, each yielding approximately the same number of edges. For the RGG, we set *r*_*c*_ = 32.6 to achieve approximately 2, 000 edges; for the ER network we selected *p* = 0.10, yielding an expected edge count of ⟨*M*⟩ = *n*(*n* − 1)*p*/2 ≈2, 000; for the SW network we chose *k* = 20, such that ⟨*M*⟩ = *nk/*2 = 2, 000. The rewiring probability *β* = 0 reproduces the initial ring lattice, whereas *β* = 1 recovers the ER limit. Analysis of degree centrality reveals that highly connected nodes cluster near the geometric center of the RGG, whereas in the ER and SW networks they are dispersed irregularly throughout the domain.

**FIG. 5.**
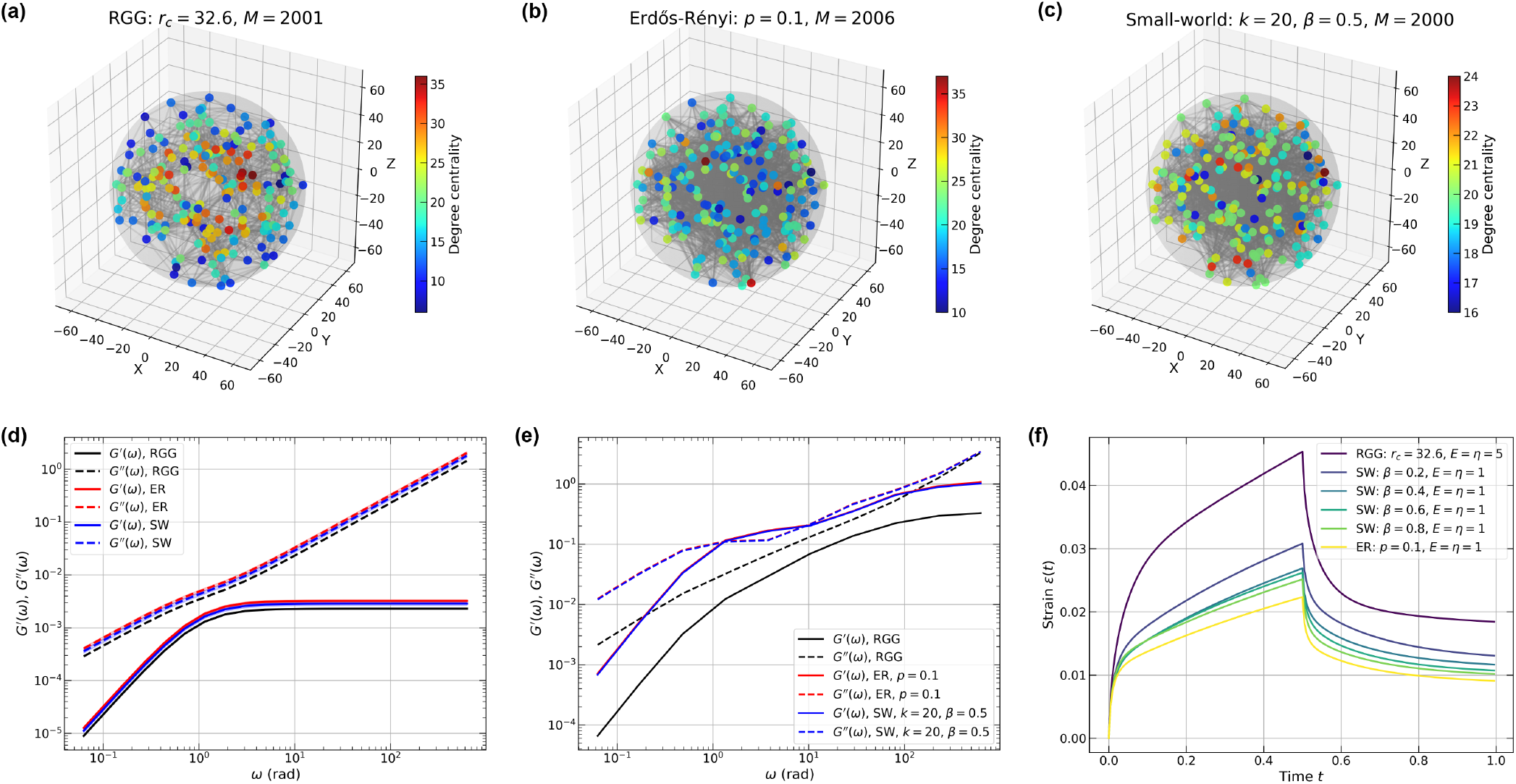
The viscoelastic responses of a random Stokes-Maxwell network with various densities. Construction of random graphs with 200 nodes inside a spherical domain with radius *R* = 63.0 and various connectivity models: (a) Random geometric graph with *R*_cut_ = 32.6; (b) Erdős-Rényi model with *p* = 0.1; (c) Small-world graph with *k* = 20, *β* = 0.5. (d) Dimensionless dynamic moduli calculated using Jeffreys model. RGG is calculated using the network in (a); ER and SW networks are each averaged over 5 samples. Shaded regions indicate standard errors. Parameters: *E* = *η* = *µ* = 1. (e) Dimensionless dynamic moduli calculated using the Stokes-Maxwell model using the same set of nodal positions but various network types. Black solid (dashed) line represents *G*′ (*G*″) for RGG; red lines: ER network; blue lines: SW network with *β* = 0.5; green lines: SW network with *β* = 0.1. Parameters: *E* = *η* = *µ* = 1 (f) Strain responses from creep tests with *σ*_0_ = 1 and *t*_1_ = 0.5 for three types of networks. RGG: *E* = *η* = 5, *µ* = 1; SW: *E* = *η* = *µ* = 1, *k* = 20, *β* = 0.2, 0.4, 0.6, 0.8; and ER: *E* = *η* = *µ* = 1, *p* = 0.1.

For comparable edge densities in the Jeffreys model, the storage and loss moduli for the RGG, ER, and SW networks collapse onto a single set of curves (Fig. 5d). Because ER and SW constructions place long- and short-range edges with equal probability, their edge-length distributions differ fundamentally from that of the RGG. These differences should engender differences in dynamic moduli, and this becomes obvious when hydrodynamic interactions are included via the Stokes-Maxwell formulation. The structural differences produce pronounced deviations in the RGG moduli, while the ER and SW results remain nearly indistinguishable across the frequency range (Fig.5e). However, the apparent similarity in dynamic moduli for ER and SW networks does not translate to similar responses in creep tests. When a constant total stress/force is applied to the same set of driven nodes in each network, the strain curves reveal significant differences as shown in Fig.5f. As the rewiring probability *β* increases from 0.2 to 0.8 in SW networks, the maximum strain progressively decreases. The ER network, which corresponds to the SW network with *β* = 1, exhibits the lowest total strain. These results indicate that SW networks possess a tunable response that is achieved by dialing the rewiring probability that interpolates between the responses of RGGs and ER networks.

Next, we developed geometric ER (GER) and geometric SW (GSW) networks to explore the effects of distance-dependent connectivity for structured graphs. Edge formation between pairs of nodes in the GER follows a distance-dependent probability *p*(*d*_*ij*_) = *p*_0_*f* (*d*_*ij*_), where *p*_0_ is the base connection probability, *d*_*ij*_ is the Euclidean distance between nodes *i* and *j*, and *f* (*d*) is a decay function that modulates connectivity based on spatial separation. We employed an exponential decay function *f* (*d*) = exp(−*d/λ*), where *λ* = *d*_max_*/α* controls the characteristic decay length, with *d*_max_ being the maximum possible inter-node distance in the network and *α* being a dimensionless decay parameter. For the results presented here, we set *λ* = *R*. This formulation ensures that spatially proximal nodes have higher connection probabilities while the stochastic nature of edge formation is preserved.

For the GSW networks, we modified the Watts-Strogatz algorithm to incorporate spatial structure: nodes are initially connected to their *k* nearest spatial neighbors, creating a spatially clustered initial topology similar to that of RGGs, followed by edge rewiring with probability *β* where new connections are formed randomly across the entire network. This approach preserves the small-world characteristics of high clustering and short path lengths while grounding the initial connectivity pattern in spatial proximity. The GSW construction differs from the original SW network in the total number of edges. Whereas each node in the SW network has exactly *k* edges before rewiring, the nearest-neighbor relationships in GSW networks are not necessarily symmetric, yielding a lower bound of *M* ≥ *nk/*2. Both geometric variants maintain the same uniform spatial node distribution as the other network types, enabling systematic comparison of how different distance-connectivity relationships affect network properties and dynamics.

When evaluating the dynamic moduli of GER versus GSW Stokes-Maxwell networks, we observe that the GER network with *p* = 0.26 and the GSW network with *β* = 1 have comparable and maximal dynamic moduli (Fig. 6c). As *β* decreases from 1 to 0.5 and then to 0, the characteristic connectivity approaches that of a RGG, thus resulting in decreased dynamic moduli, except for the converging *G*″ values at high frequencies. This transition is also evident in creep tests (Fig. 6d). The GSW network with *β* = 0 exhibits the largest maximum deformation due to its RGG-like properties; as *β* increases, the GSW network introduces randomness that approaches GER behavior, with strain curves progressively converging.

**FIG. 6.**
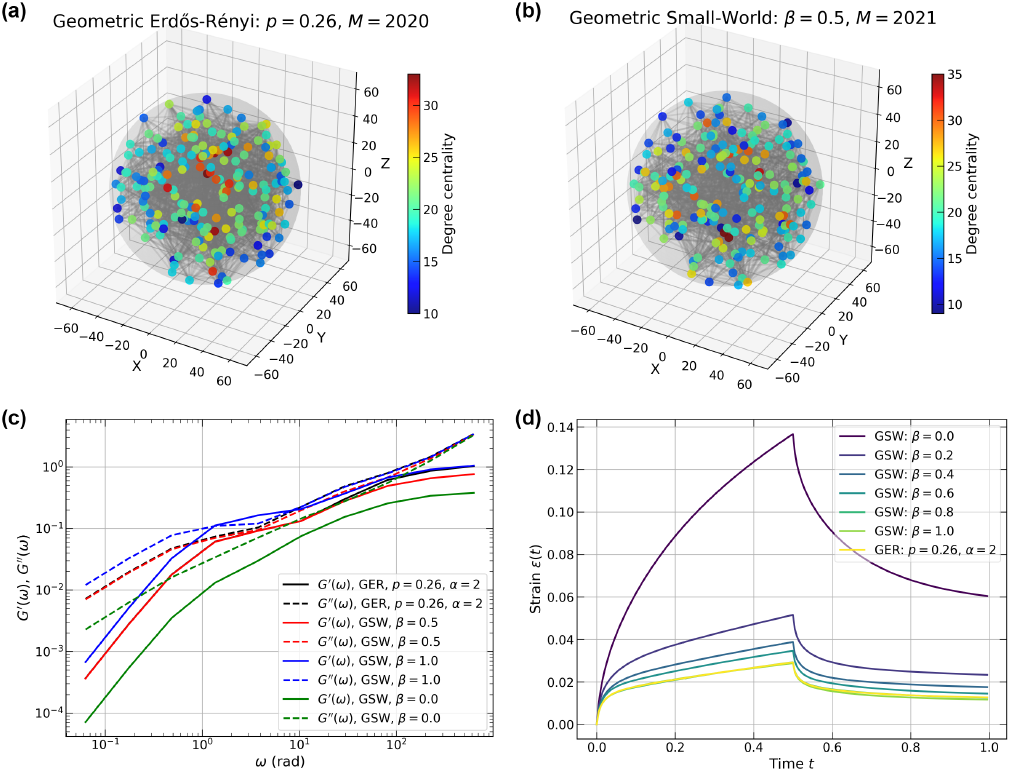
Construction of random graphs with 200 nodes inside a spherical domain with radius *R* = 63.0 and various connectivity models: (a) Geometric Erdős-Rényi model with *p* = 0.26, *λ* = *R, α* = 2; (b) Geometric small-world graph with *k* = 17, *β* = 0.5. (c) dynamic moduli calculated from Stokes-Maxwell model. (d) Strain responses from creep tests with *σ*_0_ = 1 and *t*_1_ = 0.5. GSW networks with *E* = *η* = *µ* = 1, *k* = 17, *β* = 0, 0.2, 0.4, 0.6, 0.8, 1.0.

Overall, our results show that when distance-dependent connectivity is incorporated, the approach used to generate GSW networks provides a means to explore various physically relevant scenarios through minor modifications of conventional, spatially independent ER and SW networks. GSW networks are directly relevant to the internal organization of condensates as has been shown in recent computations [70, 96, 99] and experiments [22, 25].

### E. Networks derived from coarse-grained simulations of biomolecular condensates

Analysis of condensate structures generated by coarse-grained and all-atom simulations have revealed network-like organizations of molecules [51, 57, 70, 80, 83, 96–99]. IDPs within dense phases have been shown to have high clustering coefficients [70, 96], with individual molecules making multiple contacts with several other molecules simultaneously [50, 51, 70, 80, 97, 98]. These contacts or reversible crosslinks make and break several times over the lengths of the simulations. In some cases, the high degree of clustering is reminiscent of small-world-like structures [70, 80, 96, 99].

Here, we deploy a workflow for adapting and deploying computational rheometry to quantify the viscoelastic responses of condensates that are generated using coarse-grained simulations. The workflow involves the following steps: (1) Extract the graph that best describes the network organization of molecules within dense phases. (2) Deploy the network to query how complex shear moduli vary as a function of parameter sweeps across the *E*-*η* space while keeping *µ* fixed. These parameter sweeps are best performed using the Jeffreys model to generate a diagram of dynamical states. (3) Finally, perform creep tests using the Stokes-Maxwell model for regions of the *E*-*η* space where hydrodynamic effects are important.

Different methods of coarse-graining have been brought to bear in simulations of phase separation to generate structural descriptions of condensates formed by IDPs. A popular approach is to use a single bead per amino acid residue. We first asked if different types of coarse-grained simulations, each using a single bead per residue and different potentials or sampling approaches, would generate consistent descriptions of network structures within condensates. We compared the equilibrium network structures generated using three different simulation paradigms [58, 64, 65, 68, 70, 99]. The system of interest is the well-studied prion-like low complexity domain of the protein hnRNP-A1 (A1-LCD) [10, 11]. The simulations performed using three different models were analyzed by representing each protein inside the largest cluster as a node positioned at its center of mass, with network edges modeling interactions between neighboring proteins.

To put different coarse-grained representations on an equal footing, we imposed a cutoff distance 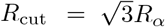, where *R*_*α*_ is the average distance between adjacent *α*-carbon atoms. When any residue from one molecule makes contact with any residue from another macromolecule within this cutoff, we establish an edge between the corresponding nodes. Details of the different simulations and the code for network conversions may be found in Appendices H-J.

Fig. 7a and 7b show representative snapshots of equilibrium configurations of two-phase systems obtained from simulations of 200 A1-LCD molecules at ~300 K and ~280 K. These snapshots were generated using three different coarse-grained simulation engines. They include a lattice-based approach (LaSSI) [70] and two off-lattice approaches namely, Mpipi-Recharged [76], and CALVADOS [65, 68, 73]. From the coarse-grained simulations, we extract networks that define condensate interiors (Fig. 7c).

**FIG. 7.**
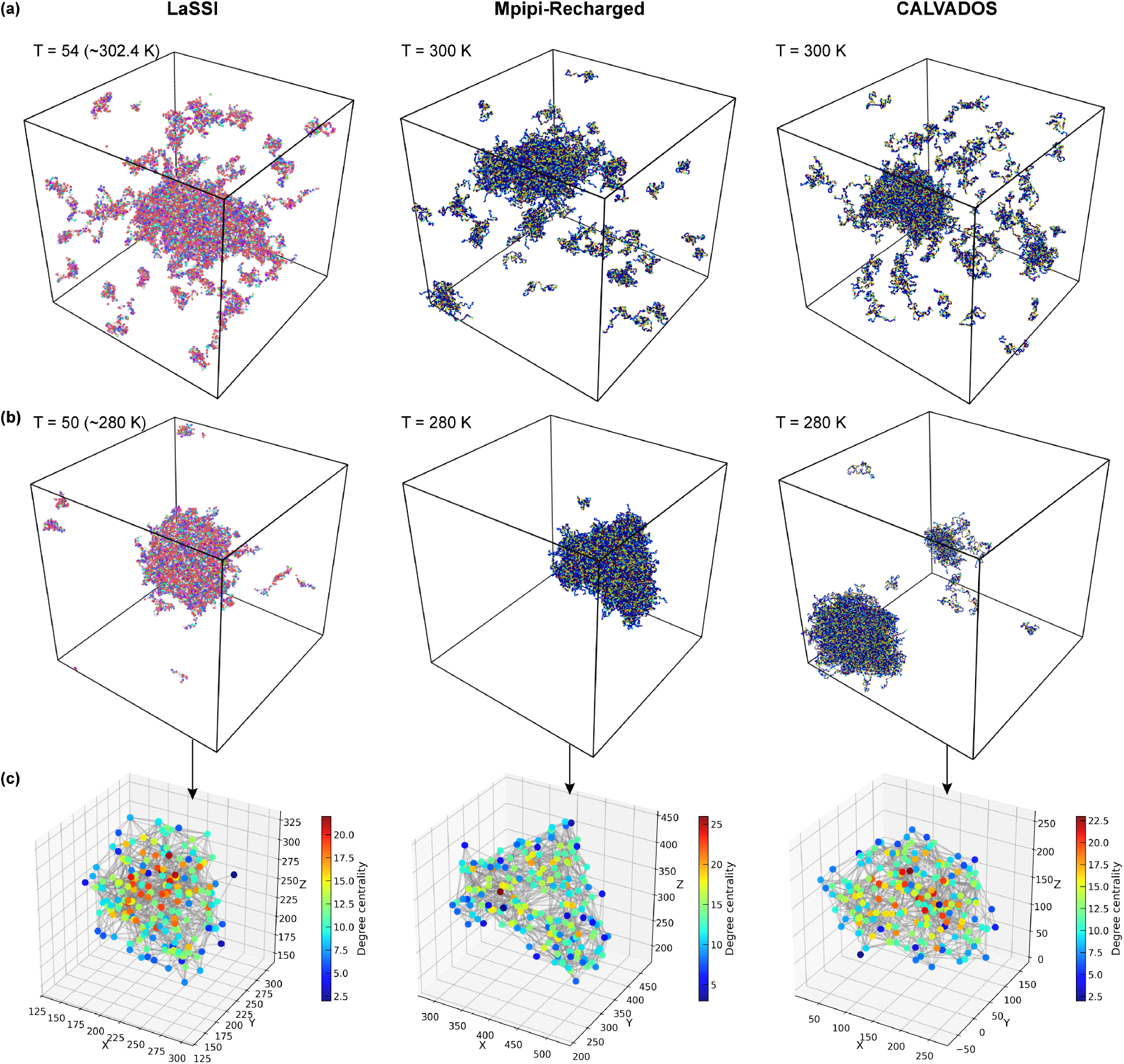
Viscoelastic networks extracted from the coarse-grained molecular simulations. (a) Snapshots from three representative coarse-grained molecular simulation engines of 200 A1-LCD proteins at ~ 300 K. (b) Snapshots at ~ 280 K. (c) A spatial network extracted from LaSSI at ~ 280 K.

The networks extracted from all three simulations exhibit small-world-like features, characterized by (*C/C*_rand_)*/*(*L/L*_rand_) *>* 3.0. Here, *C* is the average Watts-Strogatz clustering coefficient, *L* is the average shortest path length, and *C*_rand_ and *L*_rand_ are the corresponding quantities for the corresponding ER network [107, 108]. To put the comparisons on an equal footing and minimize finite-size artifacts, we performed simulations of the A1-LCD system in cubic boxes using 200 molecules and extracted temperature-dependent information regarding the network structures within dense phases of condensates. The comparisons were performed by quantifying the temperature dependence of the numbers of nodes (Fig. 8a) and the numbers of edges (Fig. 8b). These comparisons reveal a consistent temper-ature dependence with the numbers of nodes and edges decreasing with increasing temperature.

**FIG. 8.**
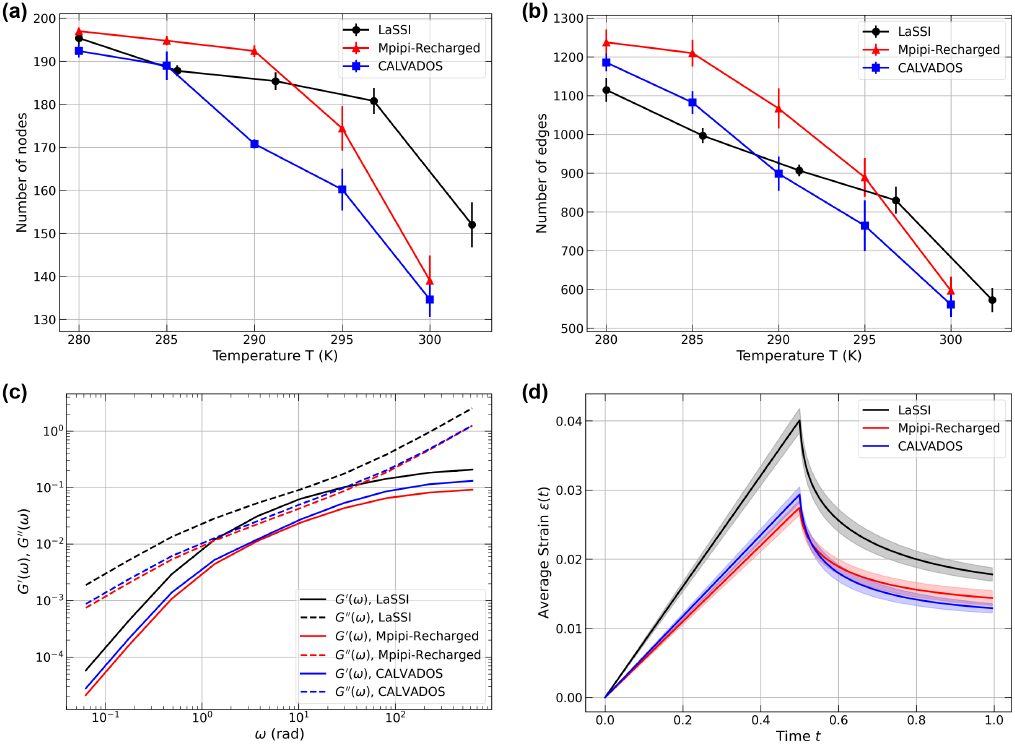
Comparison between various molecular simulation engines, i.e. LaSSI, Mpipi-Recharged, and CALVADOS. (a) Numbers of nodes and edges of the largest clusters averaged over five equilibrium snapshots. (b) Number of edges of the largest cluster averaged over five snapshots. (c) dynamic moduli of the network at *T* ~280K. (d) Creep test with constant velocity on the driven nodes.

Next, we quantified the frequency-dependent dynamic moduli using a Stokes-Maxwell representation for edges in the network. We computed the moduli by setting *E* = *η* = *µ* = 1. Snapshots for dense phases and the graphs from these snapshots were extracted for simulation temperatures of ~280 K. For this temperature, we find that all three models show similar frequency dependencies for both storage and loss moduli. For the parameters that were chosen, all graphs yield viscous moduli that are higher than storage moduli for all frequencies. We also note the absence of crossover behavior across five orders of magnitude. This is a consequence of the specific choices of the values used for the dimensionless parameters *E, η*, and *µ*.

Across the frequency spectrum, the dynamic moduli computed using the Stokes-Maxwell model for the edges remain within a factor of three of one another (Fig. 8c). The slightly larger values obtained from LaSSI-derived graphs are due to the fixed bond lengths, the discretized nature of the lattice, the formalism used to convert LaSSI simulation temperatures to Kelvin units [70] (see Appendix H), and the exact versus inexact thermostats used for Monte Carlo versus molecular dynamics simulations, respectively.

Next, we performed creep tests under constant velocity (*u*_0_ = 1) (Fig. 8d). The results show qualitatively similar behavior with minor quantitative differences. This is understandable since the LaSSI-derived network at *T* = 50 (~280 K) has comparable number of nodes but fewer edges than the other two. Overall, all three simulation methods generate mutually consistent inferences regarding the network characteristics, their temperature-dependent structures, dynamic moduli, and responses to creep tests. Our results show how the computational rheometry framework can be applied to analyze ensembles extracted from different simulation methodologies. These calculations can also be used to assess the simulated networks for equivalence or discrepancies of material properties.

### F. Evaluating the impact of boundary effects on outcomes of rheometric tests

Condensates are defined by inhomogeneous small-world-like networks [70, 71, 99]. The inhomogeneities will have an impact on mesoscale structures and dynamical responses, and this will influence the outcomes of computational rheometric tests. Additionally, there is a large gap between the lengths scales that simulations can access versus those that can be accessed in vitro. Similar discrepancies in length scale are likely to prevail between condensates that form in vitro versus in cells. These discrepancies are likely to impact the results of creep tests. Specifically, we expect there to be influences from the choices we make for the selection of boundary nodes. Therefore, we analyzed the impact of the constrained ratio on the computed dynamic moduli and the responses of networks to three different forms of creep tests. We used condensates derived from LaSSI simulations of the A1-LCD system. These simulations were performed using 2,000 instead of 200 molecules. The constrained ratio is defined as the fraction of nodes that are fixed to those than can be deformed in either oscillatory or creep tests. Fig. 9a shows the network derived from the largest cluster in a simulation containing 2, 000 A1-LCD macromolecules at a simulation temperature of *T* = 54 (~302.4K). By selecting one of the three principal axes, we define a constrained ratio *ϕ* ∈ (0, 0.5) for both the left and right boundary nodes. For oscillatory tests (performed using both Jeffreys and Stokes-Maxwell models) and creep tests (performed using the Stokes-Maxwell model), we fix the left boundary nodes while driving the right boundary nodes under various conditions.

**FIG. 9.**
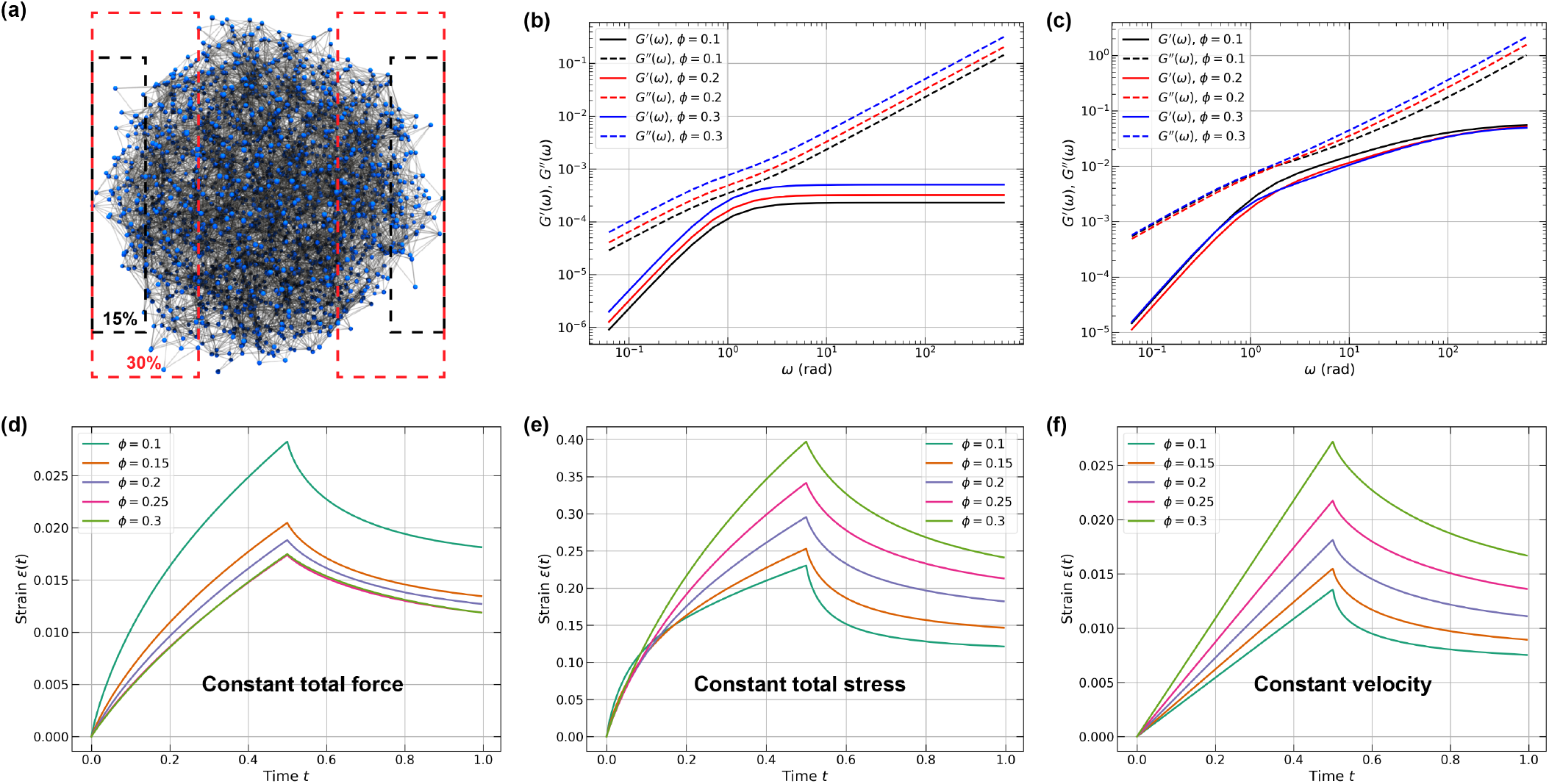
Viscoelastic responses of networks extracted from LaSSI simulations. (a) Illustration of a constrained A1-LCD network at *T* = 50 (~280K) by a variable constrained ratio in one of the three axes. Black (red) dashed rectangle: 15% (30%) of nodes in the x-direction. (b) dynamic moduli of Jeffreys networks with various constrained ratios. (c) Creep tests with various constrained ratios. (d-f) Three different creep tests on the A1-LCD WT condensate network at temperature *T* = 54 (~ 302.4K): (d) Constant total force *F*_tot_ = 1 *×* 10^4^; (e) Constant stress *σ*_0_ = 1; (f) Constant velocity *u*_0_ = 10.

Fixed boundary conditions for different rheometric tests can be imposed in two ways: (i) directly applying kinematic constraints by setting velocities to zero in all directions after solving for external forces at each time step or (ii) solving for the external forces required to maintain nodes in fixed positions. We do not observe any qualitative differences between the two methods.

The dynamic moduli of the A1-LCD network, extracted from LaSSI simulations, were first evaluated using the Jeffreys model. Again, we set *E* = *η* = *µ* = 1. As the constrained ratio increases, both storage and loss moduli shift upward (Fig. 9b). This increase can be attributed to the spatial inhomogeneities within the network: nodes within the interface, which can be identified using radial density profiles [70, 109], are more sparsely distributed and less connected than interior nodes. When the constrained ratio reaches *ϕ* = 30%, the boundary conditions encompass more densely packed and highly connected interior nodes, effectively probing the mechanical properties of the network core. Modeling the condensate as a network of Stokes-Maxwell elements reveals different behaviors with more subtle variations. While the storage moduli do not show meaningful differences among the three constrained ratios, the loss moduli exhibit a clear trend: networks with higher constrained ratios display increased loss moduli at frequencies *ω* ≳ 3.

Next, we investigated how the constrained ratio affects the results of creep tests. To ensure consistent and physically meaningful comparisons between networks of different topologies, it is essential to establish clear rheometric criteria, as networks can be deformed in numerous ways. In previous tests, we applied a constant force of magnitude *f*_0_ to each driven node, which works well for random networks with identical nodal positions in spherical domains. However, this approach becomes problematic when the total applied force varies between different network configurations. To resolve this issue, we normalized the nodal forces according to *f*_0_ = *f*_tot_*/N*_d_, where *f*_tot_ is the total force exerted on the network and *N*_d_ is the number of driven nodes. The strain response in Fig. 9d demonstrates that larger constrained ratios yield smaller maximum strains, as the force per node decreases with increasing *N*_d_, thus limiting interior node displacements. We also explored a constant-stress configuration where the nodal force is defined as *f*_0_ = *σ*_0_*A*_d_*/N*_d_. Here, *σ*_0_ represents the dimensionless total stress applied to the driven nodes, and *A*_d_ is the convex hull projection area of all driven nodes onto the plane orthogonal to the loading axis. This formulation ensures constant total stress across different configurations. The resulting strain evolution (Fig. 9e) exhibits the opposite trend compared to the constant-force case. As the constrained ratio increases, *A*_d_ increases proportionally, resulting in higher average nodal forces. Additionally, the relaxation time increases with the constrained ratio under these conditions.

Although constant-stress conditions are computationally tractable and experimentally realizable, they are less representative of typical soft matter rheometry. Standard creep tests employ either constant tensile/compressive forces or constant deformation rates. This motivated our third approach to creep tests *viz*., maintaining driven nodes at constant velocity. In this approach, we first compute the external forces required to maintain velocity *u*_0_ at the driven nodes, and then calculated the velocities of unconstrained nodes. Fig. 9f shows constant strain rates until force removal at *t* = 0.5, with larger constrained ratios producing greater maximum strains and longer relaxation times. Overall, our analysis of the A1-LCD network suggests that factors such as boundary adhesion and the choice of constant force, stress, or velocity conditions can be significantly affected by the constrained ratio, yielding quantitatively different results. Ideally, the totality of these tests will be needed both experimentally and computationally to obtain a clear understanding of the mechanical responses of condensates based on their size and the types of tests that are performed.

### G. Temperature dependence of viscoelastic responses

The phase behavior of A1-LCD is characterized by an upper critical solution temperature [10, 11, 70]. Accordingly, increasing the simulation temperature moves the system closer to the critical temperature. We performed computational rheometric tests on temperature-dependent networks derived from LaSSI simulations. Again, we set *E* = *η* = *µ* = 1. Fig. 10a and 10b show networks at simulation temperatures of *T* = 52 (~291.2 K) and *T* = 54 (~302.4 K) for a system containing 2,000 A1-LCD wild-type proteins simulated using LaSSI. As temperature increases, the interfacial regions broaden, interfacial node connectivity decreases, and the size of the largest cluster diminishes. The number of nodes and edges in the largest cluster (Fig. 10c) also follow this trend, with both quantities decreasing monotonically with temperature. Below a simulation temperature of *T* = 50, network structures remain quantitatively similar, as evidenced by consistent node counts, edge num-bers, and average degree centrality. At higher temperatures close to the critical temperature, the largest cluster forms a system-spanning network in at least one spatial direction. We therefore focused our analysis on the range of simulation temperatures from *T* = 50 to 55.

**FIG. 10.**
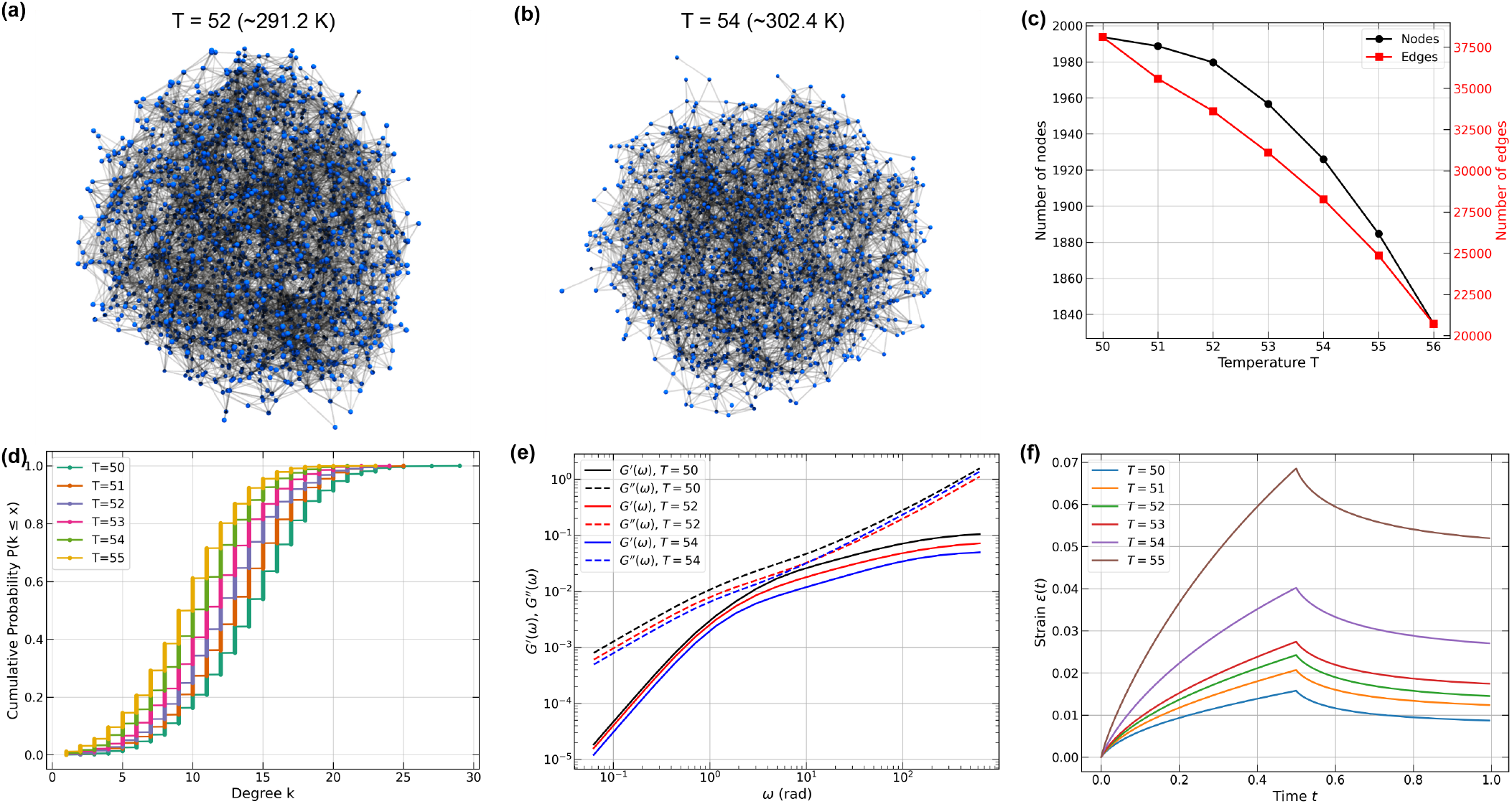
Two representative snapshots of the equilibrium structure of a 2, 000 A1-LCD system at two different simulation temperatures: (a) Simulation temperature of 52 (~ 291.2K) and (b) Simulation temperature of 54 (~302.4K). (c) Number of nodes and edges of the 2, 000 A1-LCD system at various temperatures averaged over 5 frames at equilibrium. (d) Cumulative probability function of the degree centrality at various simulation temperatures. (e) Computed dynamic moduli for A1-LCD networks at various simulation temperatures evaluated using the Stokes-Maxwell model assuming *E* = *η* = *µ* = 1. *ϕ* = 0.15. (f) Creep function of A1-LCD condensate networks at various simulation temperatures using a constant total stress.

The cumulative degree distribution shown in Fig. 10d reveals a clear leftward shift with increasing simulation temperature. This points to a systematic reduction in node connectivity. These observations demonstrate that simulation temperature affects both the volume and connectivity of the largest cluster in this closed system, suggesting corresponding changes in network viscoelasticity.

The Stokes-Maxwell model reveals systematic decreases in both storage and loss moduli as simulation temperature increases from *T* = 50 to 54 (Fig. 10e). This temperature-induced softening reflects the impact of structural changes as the critical point is approached. As interfacial regions expand and node connectivity decreases, the ability of the network to store and dissipate mechanical energy diminishes correspondingly. Creep tests under constant stress (*σ*_0_ = 0.1) further corroborate this behavior, with higher simulation temperatures producing larger deformations (Fig. 10f).

### H. Impact of changes to *E* and *η* on the responses of A1-LCD condensates

Thus far, we have investigated the responses of networks derived from LaSSI simulations by setting *E* = *η* = *µ* = 1 and probing the contributions of constrained ratios and temperature to oscillatory and creep tests. Next, we asked how the responses of an A1-LCD-like network, derived from coarse-grained LaSSI simulations, changes as we vary *E* and *η* while keeping the solvent viscosity fixed at *µ* = 1. We assume homogeneity of the elements within the inhomogeneous network by setting equivalent values of *E* and *η* for all the elements. We selected a highly connected 2, 000-chain A1-LCD network configuration drawn from ensembles obtained at a simulation temperature of *T* = 50 (~280 K). We start with a reference state wherein *E* = *η* = 10.

Symmetrical increases in *E* and *η* cause a uniform up-ward shift in *G*′ across all frequencies (Fig. 11a). As the network stiffens, the storage modulus increases across the entire frequency range. The loss modulus *G*″ follows a similar trend for *ω* ≲ 10 rad. However, for higher frequencies we observe convergence onto a universal curve. Importantly, we observe double crossover behaviors for all three sets of *E* and *η* values. For *E* = *η* = 5, the crossovers occur at frequencies of *ω* ≃ 1 and 10 rad. As we increase *E* and *η*, the gaps between the two crossover frequencies widen. While the lower crossover frequency remains nearly constant, the higher crossover shifts by an order of magnitude as *E* and *η* increase from 5 to 100.

**FIG. 11.**
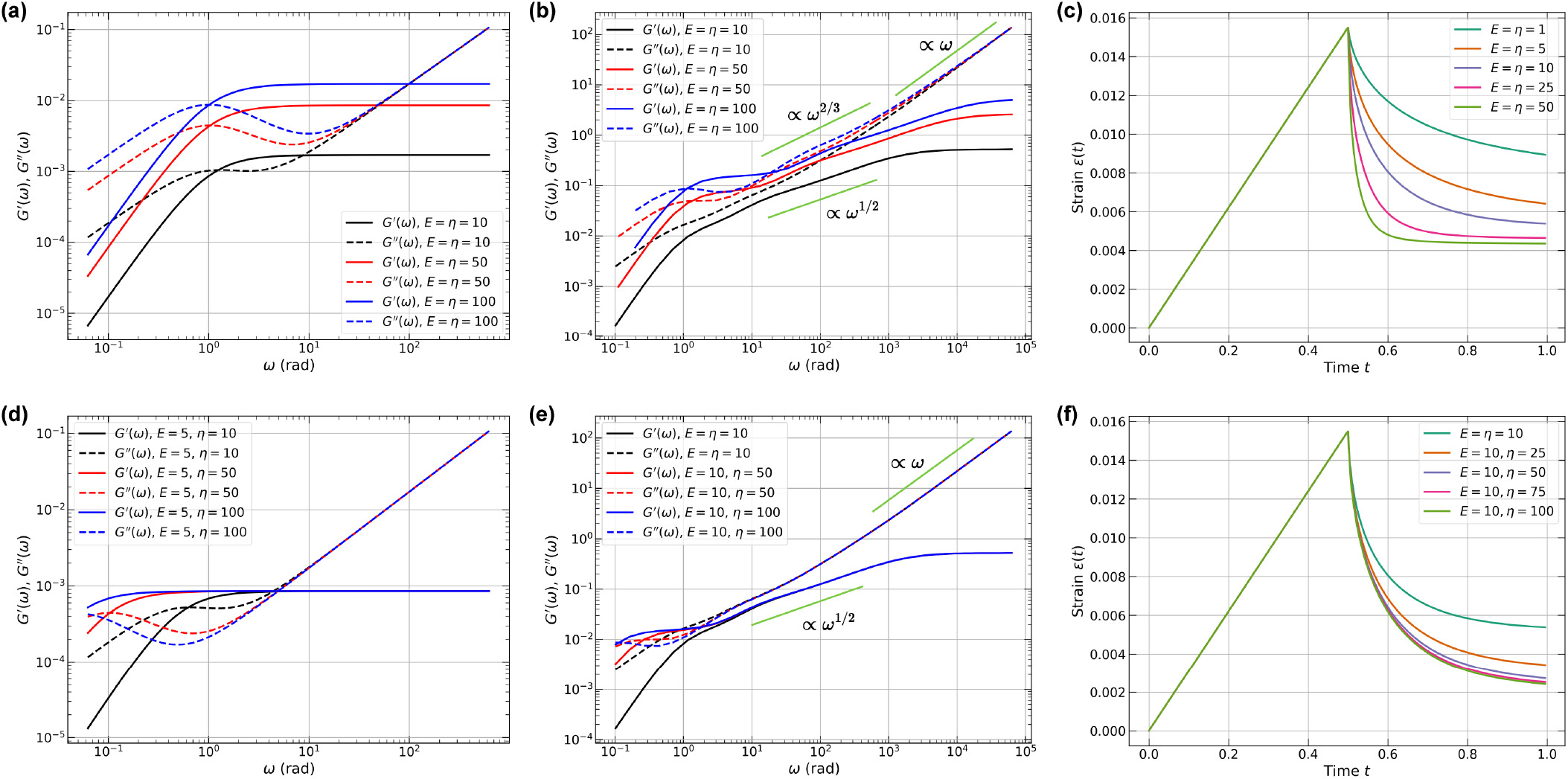
Effects of model parameters on viscoelastic response patterns on an A1-LCD condensate. (a,b) Dimensionless dynamic moduli as *E* and *η* increases with the same magnitudes using (a) the Jeffreys model and (b) the Stokes-Maxwell model. Green lines: scaling slopes as guides to the eye. (c) Creep test with constant velocity for equal value of *E* and *η*. Constant velocity *u* = 10. (d,e) Dimensionless dynamic moduli as *E* remains constant and *η* increases using (d) the Jeffreys model and (e) the Stokes-Maxwell model. Green lines: scaling slopes as guides to the eye. (f) Creep test with constant velocity for constant value of *E* = 10 and varying *η*.

The presence of a double crossover is pertinent in light of the results of Galvanetto et al., [51]. For the most well-studied variant of their systems, they estimated a single crossover frequency of ~1 MHz using the reciprocal of the measured single chain reconfiguration time, which is in the microsecond timescale. However, the reconfiguration time for a single chain is not the same as the reconfiguration time of a network [83]. The network reconfiguration time, which can be extracted from measured complex shear moduli by solving an inverse problem [83], is likely to be on the 0.1 - 10 s range. This estimate is concordant with the measured zero shear viscosities for different condensates [8, 26, 28, 36, 37, 45, 51, 52, 110]. The characteristic timescale for network reconfiguration and its convolution with single chain dynamics will generate at least one additional crossover at low frequencies (*ω*_low_). Detecting the presence of one or more crossover frequencies requires the measurement of the full frequency-dependent dynamic moduli because this probes responses on intermediate length and timescales [26, 28, 34–36, 41, 111] that go beyond measurements of dynamics of individual chains and zero shear viscosity of condensates.

The Stokes-Maxwell formalism reveals markedly dif-ferent behaviors as *E* and *η* are varied symmetrically (Fig. 11b). Double crossover behavior emerges at higher values of *E* and *η* when compared to the Jeffreys model, though still within 1 *< ω <* 100 rad. Unlike the plateauing of *G*′ observed for the Jeffreys model, *G*′ in the Stokes–Maxwell model exhibits an intermediate scaling regime where *G*′∝ *ω*^1*/*2^, before plateauing at much higher frequencies. The loss modulus displays even more nuanced behaviors: there is an intermediate regime where *G*″∝ *ω*^2*/*3^. This behavior was recently observed in DLS experiments performed by Fisher and Obermeyer for coacervates formed by acidic peptides complexed with a cationic fluorescent protein [41]. Their work provides an unprecedented estimation of dynamic moduli across nearly six orders of magnitude and it appears to be the first report of the double crossover behavior and the *ω*^2*/*3^ dependence of *G*″ for condensates/coacervates. In the high frequency regime, we observe a *G*″∝ *ω* scaling, that coincides with expectations for the Jeffreys model.

Next, we performed creep tests and queried the responses to symmetric changes in *E* and *η* while keeping *µ* fixed. These tests were performed under constant velocity conditions. The creep tests, performed for A1-LCD graphs, modeled as networks of Stokes-Maxwell elements, reveal the time-dependent consequences of symmetric changes to *E* and *η* (Fig. 11c). During loading, all networks exhibit identical strain evolution due to the imposed kinematics. However, upon removal of the constraints, relaxation behavior depends critically on the material parameters. As *E* and *η* increase, the relaxation time decreases dramatically, with networks showing negligible relaxation for *E* = *η* ≥ 50. This points to time-invariant residual strain.

The effects of changes to *E* and *η* may be viewed as tests of the contributions from dynamical arrest or physical aging responses that have been observed for a variety of condensates [23, 26, 28, 32, 34, 35, 46, 76, 112–119]. There are two sets of timescales and two types of processes to consider when discussing aging. These are *t*_age_ and *t*_obs_. Here, *t*_age_ refers to the physical age of the condensate with *t*_age_ = 0 being the time at which the condensate first forms; *t*_obs_ refers to the timescale that is spanned by a rheometric measurement (cf., the frequency range along the abscissae of each of the panels in Fig. 11). Aging can come about due to increased elasticity, increased viscosity, or symmetric increases in both. In a recent computational study, Biswas and Potoyan [120] investigated the effects of changing the lifetimes of physical crosslinks between specific pairs of residue types referred to as stickers. They noted that short-lived stickers may lead to a Maxwell fluid behavior, while longer-lived, irreversibly cross-linked stickers may result in solid-like properties, consistent with the Kelvin-Voigt model [120].

We investigated how varying *η* while keeping *E* and *µ* fixed affects the dynamical and mechanical responses of A1-LCD condensates derived from LaSSI simulations. We first used the Jeffreys model and increased *η* from 10 to 100 while maintaining *E* = 10. This transforms the network from balanced elasticity and viscosity of the elements that are in series to a viscosity-dominated regime. As predicted by single-element analysis (cf. Fig. 2d), this pathway produces a characteristic leftward shift of the lower crossover frequency (*ω*_low_) while preserving the upper crossover position (*ω*_high_). Remarkably, both moduli become independent of *η* for *ω* ≳ 5 rad, indicating that the high-frequency responses are governed entirely by the elastic components (Fig. 11d).

The Stokes-Maxwell formalism reveals additional complexity in response to changes in viscosity that occur without changes to elasticity (Fig. 11e). While the high-frequency convergence remains (*ω* ≳ 5 rad), an extended intermediate regime emerges with anomalous scaling whereby *G*′∝ *ω*^1*/*2^. This regime spans nearly two orders of magnitude in frequency. The *G*′∝ *ω*^1*/*2^ scaling is absent in the Jeffreys model. Thus, hydrodynamic interactions can impact the elastic responses and how viscous dissipation manifests at intermediate timescales. The crossover regions for both Jeffreys and Stokes-Maxwell models, 0.1 *< ω <* 10 rad, occur at a lower frequency regime than the case where *E* and *η* change symmetrically.

Next, we investigated creep relaxation while fixing *E* and varying *η*. Increasing the viscosity of each of the elements in the network produces progressively slower but persistent relaxation (Fig. 11f). This contrasts with the abrupt relaxation and arrest observed in symmetric changes to *E* and *η* (cf., Fig. 11c). This distinction has profound implications for age-dependent mechanical responses of condensates: Symmetric changes to *E* and *η* create effectively permanent deformations. In contrast, fixing *E* while changing *η*, a scenario that might imply the absence of structural changes despite the onset of dynamical arrest, maintains the capacity for complete recovery given sufficient time. These results highlight the importance of performing creep tests while separately probing structural transitions as well as deploying temperature-jump studies [28, 32]. It also highlights the importance, in computations and experiments, of deploying tests that probe the effects of changing the lifetimes of physical crosslinks, as reported by Biswas and Potoyan [120], and probing the effects of structural changes as prototyped by Collepardo and colleagues [76, 115, 121].

## IV. DISCUSSION

The Rouse model for viscoelasiticity of polymer solutions may be viewed as being a generalized Maxwell model that comprises Maxwell elements in parallel, with each element being embedded in a viscous, incompressible fluid [92]. The Rouse model ignores the contributions of hydrodynamic interactions. The Zimm model, which is a generalization of the Rouse model, considers individual Maxwell elements being influenced by hydrodynamic interactions [93]. We derived two limiting scenarios for hydrodynamic interactions. In the strongly screened limit, each element in the viscoelatic network may be modeled as a Jeffreys element. In the opposite, strong hydrodynamic coupling limit, the network can be modeled as Maxwell elements coupled to a Stokes fluid. Zhou has proposed that condensates are best described as viscoelastic Jeffreys fluids [82]. Our analysis suggests that condensates are likely to exist between the strong screening and the strong coupling limits. Therefore, we adopted an approach of modeling the mechanical responses for both types of networks using the computational rheometry formalism [84].

We used the regularized Stokeslet approach of Wróbel et al. [84] for modeling viscoelastic networks in a Stokes fluid. For individual Jeffreys and Stokes-Maxwell elements we probed the responses to oscillatory displacements. For the Stokes-Maxwell element, we also performed creep tests. Although we find convergent behaviors for single elements, in general, the Stokes-Maxwell model shows more complex dynamical behaviors when compared to the Jeffreys model when we consider different types of networks. The differences become significant and non-trivial when the networks have structural inhomogeneities and are defined by more than just connectivity considerations, as is the case for graphs derived from simulations that model the small-world-like internal organization and dynamics of condensates. The full range of complexities of measured dynamic moduli, as reported by Fisher and Obermeyer [41], are only evident when we use the Stokes-Maxwell model.

We presented and prototyped a workflow for deploying computational rheometry as an approach that bridges the molecular and mesoscales. This is achieved via graph-based representations of networks of molecules and using computational rheometry to model the responses of networks where each edge or node in the network is a viscoelastic element. We showed how networks can be extracted from analysis of coarse-grained simulations of condensates formed by an archetypal IDP. Similar approaches can be brought to bear to analyze all-atom simulations or other coarse-grained models that use more than one bead per residue or even single beads or rods for entire domains [57, 122–124].

We focused on the responses due to structural inhomogeneities within different types of networks. However, it is possible that there are additional inhomogeneities present in terms of the parameters themselves. Recent studies have shown that folded domains are themselves viscoelastic elements [125]. They are likely to be different types of viscoelastic elements when compared to IDPs. Hence, it is likely that the viscoelastic elements within networks are defined by element-specific values for *E* and *η*. This would entail a form of symmetry breaking that we have not considered in the current work. Even for IDPs, there likely are heterogeneities in terms of the *E* and *η* parameters for each of the elements within condensates. Interaction hierarchies defined by hydrogen bonds, hydrophobic contacts, and the spectrum of electrostatic attractions are likely to contribute differently to mechanical responses of networks. Indeed, a recent study using a polarizable forcefield showed that the polypeptide backbone is the key generator of networking and percolation within dense phases of peptide-based mimics of condensates [80]. To accommodate these hierarchies, one would need finer-grained simulation models from which the networks are derived. Different weights for different edges are likely to be a useful consideration. All-atom models are advantageous because they do not require any additional parameterization beyond the parameters specified by the molecular mechanics forcefield [50, 51, 79, 81]. The high computational cost of all-atom models may be offset by deploying implicit solvent models with solvation inhomogeneities [126] or coarse-grained models with multiple interaction sites per residue [72, 77, 127, 128]. Our framework readily accommodates such extensions through edge-specific or node-specific parameterization, enabling future studies that might capture molecular-level heterogeneity.

The key takeaway from our work is that even for uniform choices that we make for *E* and *η* values, we need an entire gamut of rheometric tests to identify the appropriate structural description, the suitable model, and parameters for the elements that best describe the mechanical responses of condensates. No single test proves to be sufficient for adjudicating how the mechanical responses of condensates are determined by the microstructures and their inherent dynamics. The choice between edge-based (Jeffreys) and node-based (Stokes-Maxwell) approaches involves trade-offs between computational efficiency and implementation flexibility. The Jeffreys model offers substantial computational advantages, as we only need to solve a linear problem in the frequency domain, making it at least an order of magnitude faster than the iterative solutions needed for the Stokes-Maxwell system. This efficiency makes the Jeffreys approach particularly suitable for analysis of experimental data via parameter sweeps. This does implicitly impose the assumption of hydrodynamic screening. Importantly, the use of the Jeffreys model does not require the construction of graphs that match the sizes of condensates measured in experiments, which tend to be on the micron scale. In contrast to the Jeffreys model, the Stokes-Maxwell formalism becomes essential for querying the effects of hydrodynamic coupling. Although it carries a higher computational cost, the Stokes-Maxwell approach offers greater flexibility in boundary condition specification and naturally captures long-range hydrodynamic coupling through the fluid phase. However, numerical stability becomes challenging at low frequencies with large *E* and *η* values (e.g. Fig. 11b and Fig. 11e). This in turn requires a small *dt*, resulting in longer simulation times. Future implementations could benefit from adaptive time-stepping algorithms and parallel linear solvers such as PETSc [129] to extend the accessible parameter range and reduce computational time. In addition to numerical stability considerations, the proper use of the Stokes-Maxwell model will likely require the construction of micron scale graphs if we are to simulate true facsimiles of experimental observations. While simulations of condensates that approach or exceed the micron-scale are likely to be improbable, it should be possible to finite-size corrections [130] and finite-size scaling [131] to rescale the computed graphs to make them be suitable facsimiles of measurements.

For creep tests, we exclusively employed the Stokes-Maxwell formalism due to its versatility in handling diverse boundary conditions within a unified solver framework. Overall, the two approaches serve complementary roles: the Jeffreys model provides rapid screening and parameter estimation, while the Stokes-Maxwell formalism delivers physically complete descriptions when hydrodynamic effects cannot be ignored.

The current formalism rests on the assumption of linear viscoelasticity, which is appropriate for small deformations but insufficient for large strains. Extending to nonlinear regimes would require incorporating strain-dependent edge properties. Viscoplastic deformations such as bond rupture mechanics where edges permanently deform or break beyond critical strain thresholds can also be included to model irreversible transitions within networks. Such extensions would better capture phenomena like strain stiffening, yielding, and fracture that might be relevant in non-equilibrium deformations of condensate networks and cellular structures.

The Stokes-Maxwell formalism presented here exemplifies continuum-particle coupling methods that can be extended to more complex experimental geometries. Previous work has demonstrated the versatility of this approach for modeling micropipette aspiration tests [84], suggesting natural extensions to other rheometric configurations. Of relevance for biomolecular condensates would be the incorporation of probe particles to study their interactions with heterogeneous network structures.

Our work bridges multiple scales of description, from macromolecular networks to continuum models. At the coarser end, the Stokes-Maxwell system can be mapped onto into Oldroyd-B-type constitutive equations for viscoelastic fluids, enabling connection with continuum theories of viscoelastic phase separation that have been developed to describe polymer solutions and the dynamics of viscoelastic spinodal decomposition [132]. This connection suggests that network-based parameters could guide continuum-level descriptions, providing a systematic route for connecting molecular structure to mesoscopic and macroscopic rheology.

## ACKNOWLEDGMENTS

This work was supported by grants from the US Air Force of Scientific Research (FA9550-20-1-0241), the US National Institutes of Health (R01NS121114), the US National Science Foundation (MCB-2227268), and the St. Jude Research Collaborative on the Biology and Biophysics of RNP granules. We owe special thanks to Priya Banerjee for inspiring us to pursue computational rheometry and to Samuel Cohen for guidance during the formative stages of this work. We are grateful to Clifford Brangwynne, Tuomas Knowles, Tanja Mittag, Michael Rosen, and Andrea Soranno for helpful discussions.

## AUTHOR CONTRIBUTIONS

R.Z. and R.V.P. conceptualized the adaptation of computational rheometry and designed the workflow; R.Z. adapted the framework and developed the code; R.Z., G.M., S.G., and R.V.P. analyzed the results; R.Z. and R.V.P. wrote the manuscript. All authors read and edited the manuscript. R.V.P. secured funding.

## CODE AVAILABILITY

The code for reproducing the results will be made available at GitHub (https://github.com/Pappulab/Viscoelastic_network).

## Appendix A: Hydrodynamic screening length of a viscoelastic network in Stokes flow

Following the work by Levine and Lubensky [90, 91], we consider a viscoelastic network characterized by a displacement field **u**(**r**, *t*) when it is immersed in a viscous solvent with velocity field **v**(**r**, *t*). The coupled dynamics are governed by momentum balance equations:

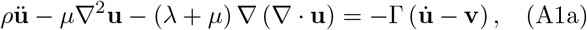

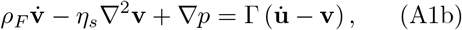

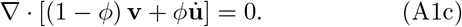

The network is assumed to be dilute (*ϕ* ≪ 1) and incompressible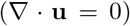. We also assume the limit of low Reynolds numbers (Re = *ρ*_*F*_ *V L/η*_*s*_ ≪1) and regime of linear viscoelasticity, such that network response of the network to small amplitude oscillatory shear is characterized by the complex shear modulus *G*^*^(*ω*) = *G*′(*ω*) + *iG*″(*ω*). Rearranging Eqs. A1a-A1c and applying a Fourier transform we obtain,

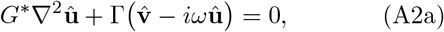

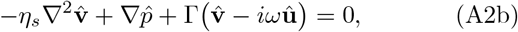

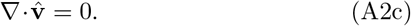

The spatial Fourier transform using Δ^2^ → − *k*^2^ and Δ→ *i***k** yields,

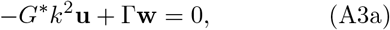

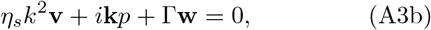

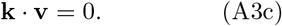

Here, we define the relative velocity field **w** ≡ **v** −*iω***u**. Since we focus on the shear response, we project the fluid momentum balance onto the transverse projector **P**(**k**) = **I** − **kk**^⊤^*/k*^2^. As a result, the pressure term in Eq. A3b vanishes and **P**(*i***k***p*) = 0. From Eq. A3a, we solve for **u**:

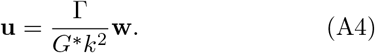

Since **v** = **w** + *iω***u**,

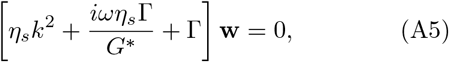

which yields the solution for **w** as:

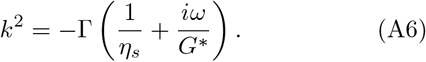

We define a frequency-dependent screening parameter *κ*^2^(*ω*) = −*k*^2^(*ω*) and return to real space. The relative velocity **w**(**r**, *ω*) obeys the Helmholtz equation for each Cartesian component:

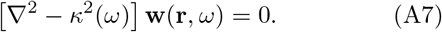

Assuming spatial isotropy and radial symmetry about a localized forcing center, we consider a scalar component *ψ*(*r*) of the form:

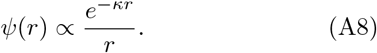

For a complex *κ* = *κ*′ + *iκ*″, the magnitude of any component of **w** decays as

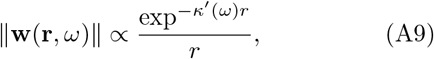

where *κ*′ *>* 0. We thus define a hydrodynamic screening length *ξ*(*ω*) by:

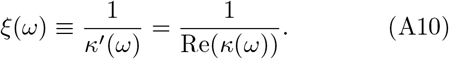

To express *ξ*(*ω*) explicitly, we write:

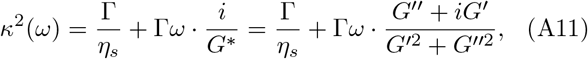

so that:

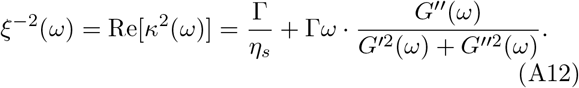

Finally, we define a dimensionless coupling parameter that quantifies the relative strength of hydrodynamic screening at frequency *ω*:

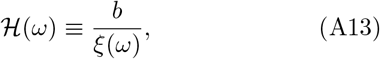

where *b* is the characteristic mesh size of the network. If the network is purely elastic, i.e., *G*″ = 0, the expression can be simplified to to 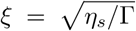 and subsequently 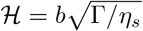, which is frequency-independent.

We can make further estimation of the friction coefficient Γ. Consider a node (bead) that moves relative to background fluid at a relative velocity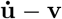. The drag force it experiences is 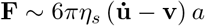,where *a* is the characteristic size of the node. We approximate the network cell volume as *b*^3^, where *b* is the characteristic mesh size. From Eqns. A1, we let the force per unit volume be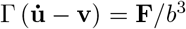, leading to Γ ~ 6*πη*_*s*_*a/b*^3^. In the work by Levine and Lubensky [91], a single length scale *b* is used to characterize the network, reaching a equivalent Γ ~ *η*_*s*_*/b*^2^ expression.

In the context of biomolecular condensates, we can estimate the typical range of the coupling parameter ℋ using experimentally relevant values. The solvent viscosity *η*_*s*_ ranges from 1.0 mPa· s for water to 100.0 mPa· s for various biofluids [133]. Molecular network mesh sizes have been reported between 5 and 20 nm [29, 49]. For physically relevant frequencies of 0.1 to 100 Hz, experimental measurements yield dynamic moduli typically spanning 0.01 to 100 Pa [26, 28]. This yields an estimate of 0.1 *<* ℋ *<* 2, indicating that biomolecular condensates operate in an intermediate regime where neither limit (ℋ ≫ 1 or ℋ ≪ 1) fits perfectly. Consequently, both local network elasticity and long-range hydrodynamic interactions contribute significantly to the mechanical response. This analysis suggests that a complete description of condensate rheology requires models that capture both effects simultaneously. Accordingly, as noted in the main text, we analyze every network of interest using the Jeffreys and Stokes-Maxwell models identifying shared and discrepant behaviors for both models.

Finally, we defined the mesh using the center of mass of each chain, which may overestimate the actual mesh size. However, this approximation enables coarsegraining from molecular structure to continuum representation.

## Appendix B: Non-dimensionalization

The system is non-dimensionalized following the work of Wróbel et al. [84]. All variables are cast into a dimensionless form by three reference scales: *L* (length), *U* (velocity) and *µ*_0_ (dynamic viscosity). For example, the dimensional coordinates and velocities 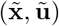 are mapped to 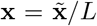 and 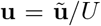, respectively. The time scale is *T* = *L/U*, so that 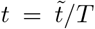. Here, a tilde denotes dimensional quantities; symbols without tilde are dimensionless.

The parameters for describing material properties used in this work in Sec. II are scaled as follows: *E* = 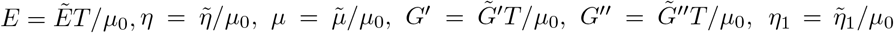 and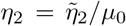. We choose 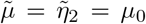 so that the dimensionless solvent (dashpot) viscosities appear as *µ* = *η*_2_ ≡ 1. Angular frequency is scaled by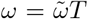, and the dynamic moduli are plotted against *ω* in radians.

Throughout the manuscript we choose the timescale such that the time taken for a node to displace a dimensionless distance Δ*x* = 1 with velocity *u* = 1 will take exactly Δ*t* = 1. For the molecular network in this work, the characteristic length is set to 1 Å for easier comparison with lattice and random networks. To non-dimensionalize a physical system, for example, we can take *L* to be the molecular mesh size, *U* to be the characteristic bead velocity (used in optical tweezers), and *µ*_0_ the viscosity of water. All lengths, velocities and viscosities are then expressed as ratios with respect to *L, U*, and *µ*_0_, respectively.

## Appendix C: Frequency-domain formulation for a Jeffreys network

We consider an undirected network 𝒢 = (𝒱, ℰ) embedded in ℝ^3^. Symbols used in the model are defined in Table I. For every node *i* ∈ 𝒱, let **r**_*i*_ ∈ ℝ^3^ be its reference position and 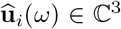 the complex amplitude of its harmonic displacement at angular frequency *ω*. For an edge *e* = (*i, j*) ∈ ℰ we define

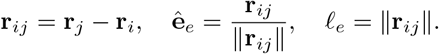

**TABLE I.**
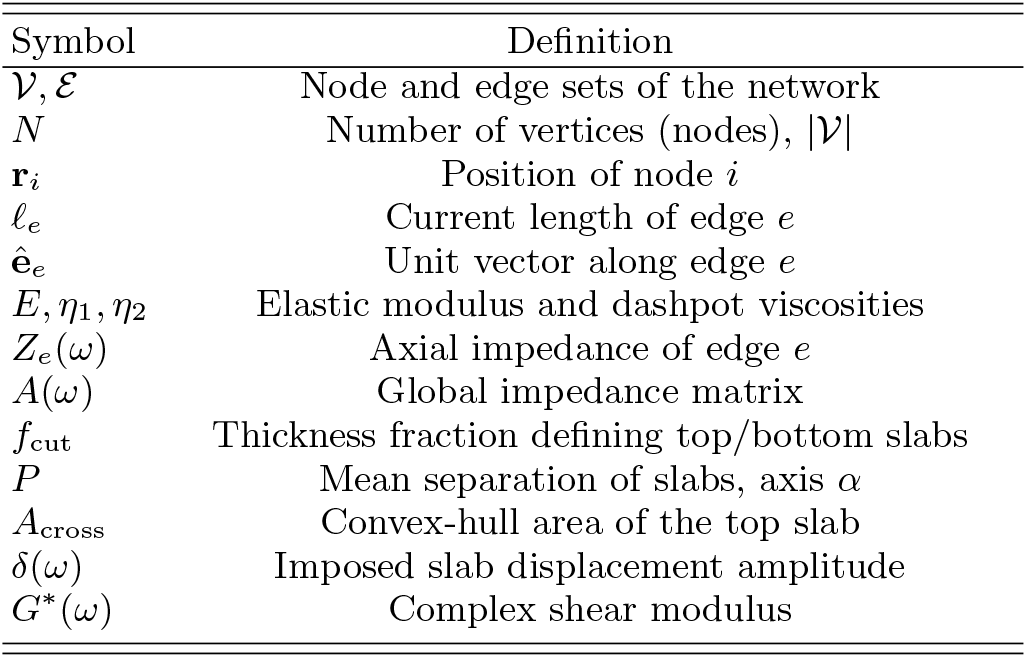
A table for symbols used in Appendix C.

We compute stress-strain relationships based on the original cross-sectional area and length of the element. The strain is therefore referred to as the engineering strain and the stress is the engineering stress. In contrast, true stress-strain relationships use the instantaneous cross-sectional area and length. The small engineering strain in edge *e* is,

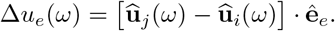

Each edge is a one-dimensional Jeffreys element whose material parameters *E, η*_1_, *η*_2_ are assumed uniform in space. With a unit cross-section of the bar *A*_*e*_ = 1, the one-dimensional constitutive coefficients are *k*_*e*_ = *E/*𝓁_*e*_, *η*_1,*e*_ = *η*_1_*/*𝓁_*e*_, and *η*_2,*e*_ = *η*_2_*/*𝓁_*e*_. The resulting complex axial impedance is

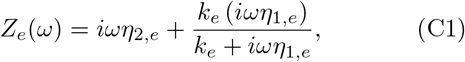

and the axial force amplitude in the edge is 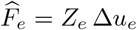.

For edge *e* = (*i, j*) form the 3 3 tensor 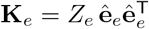acts on the displacement difference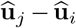. Denoting by 𝒫_*e*_ the Boolean (3*N*) × (3*N*) projector that inserts +**K**_*e*_ into the diagonal blocks (*i, i*) and (*j, j*) and −**K**_*e*_ into the off-diagonal blocks (*i, j*) and (*j, i*):

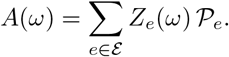

We choose one Cartesian axis *α* (*α* = 0, 1, 2 ≡ *x, y, z*). Nodes whose *α*-coordinate lies in the lowest fraction *f*_cut_ of the span form the *bottom* boundary *B* and are fixed. Nodes in the highest *f*_cut_ fraction form the *top* boundary *T* and are prescribed a sinusoidal displacement

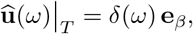

where *β* ≠ *α* is the loading direction (e.g. simple shear in *y* if *α* = *x*). All remaining nodes constitute the interior set *I*. Partition *A*(*ω*) accordingly:

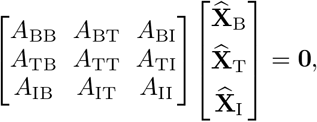

with boundary conditions 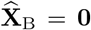 and 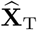 prescribed. The only unknown displacements (the interior DOFs) 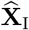 can be solved by

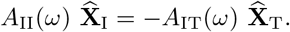

Here we note that a small (*ϵ* = 1 ×10^−12^) regularization parameter is added to the diagonal entries of *A*_II_ to stabilize the solver. After obtaining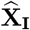, we can then construct the full displacement vector 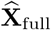

The top-slab reaction force is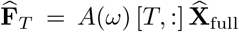. By summing its *β*-components, we have the net shear force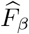. Let *A*_cross_ be the convex-hull area of the top slab projected onto the plane perpendicular to axis *α* and *P* the mean *α*-separation of the two slabs. The strain amplitude is thus *γ* = *δ/P*. Finally, the complex shear modulus is evaluated as follows:

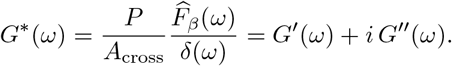

## Appendix D: Numerical methods for Stokes-Maxwell networks

Throughout we treat the discrete network and the surrounding solvent as a fully coupled Stokes-flow problem. The *N* network nodes are located at **x**_*i*_(*t*) ∈ℝ^3^; every internal node is linked to its neighbors by a linear Maxwell element. The solvent is incompressible with dynamic viscosity *µ* and is described by the method of regularized Stokeslets. All quantities in the code are already non-dimensional; here we retain the same symbols. The numerical methods used are consistent with the existing models in the literature [84, 103, 134].

The regularization function in Eq. 5 is defined as [84, 103]

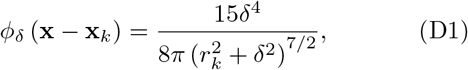

where the parameter *δ* controls the width of the function.

### 1. Creep and related tests

The creep test, as originally defined, is a mechanical test that can be used to evaluate how a material deforms over time under constant stress. In Sec. III F, we discussed the alternative forms of such tests whereby we evaluate deformation under the influence of constant stress, constant total force, and constant velocity, respectively. Here we lay out the numerical method used for the application of uniform normal stress.

For a link connecting nodes *i* and *j* we denote the current length *r*_*ij*_ = ∥**x**_*j*_ −**x**_*i*_∥, the time-dependent Maxwell rest length 𝓁_*ij*_(*t*) and the initial rest length 𝓁_*ij*,0_. The elastic (Hookean) force acting on node *i* is

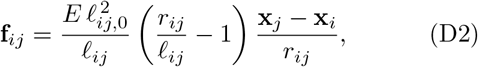

where *E* is the one–dimensional elastic modulus. When a dashpot of viscosity *η* is in series (Maxwell link) the rest length evolves as

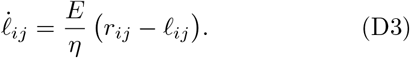

Let 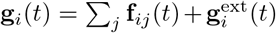 be the net force on node *i*. The associated velocity is obtained from the regularized Stokeslet mobility as,

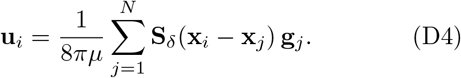

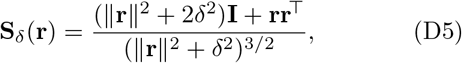

The regularization parameter is *δ* = 𝓁_*ij*,0_*/*(4*π*). In the code, we compute this once and stores the tensor **S**_*ij*_. For simplicity, we choose *δ* = 1*/*(4*π*) for all cases. Care is taken to ensure that 𝓁_*ij*,0_ ≫ *δ*.

Nodes whose *x*-coordinate lies in the leftmost *ϕ* fraction of the network are fixed; nodes in the rightmost *ϕ*-slice are driven. A uniform normal stress *σ*_0_ is imposed for *t < t*_1_ by applying

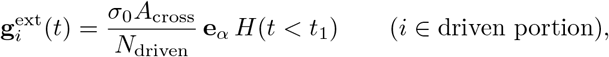

where *A*_cross_ is the instantaneous cross–sectional area and **e**_*α*_ the load axis (*α* = *x, y, z*). Fixed nodes satisfy **u**_*i*_ = **0**. At every step the unknown Lagrange multipliers, which represent reaction tractions on fixed nodes, are determined by solving 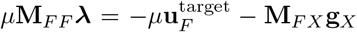 with **M**_*F F*_ the mobility subblock among the *N*_*F*_ fixed nodes and **M**_*F X*_ the block from (D4). The ODE state vector is **y** = (𝓁_*ij*_, **x**_1_, …, **x**_*N*_) and it is integrated with `dopri5` (SciPy package with default tolerance).

The engineering strain along the loading axis is calculated using Eq. 10. The code records *ε*(*t*), the strains per-link, the node velocities, and the three–dimensional velocity field using a user-specified Cartesian grid.

### 2. Small-amplitude oscillatory shear test

For a given network, the full 3*N*× 3*N* mobility matrix **M** from (D4) is assembled once. We partition the node indices into ℱ (fixed) and ℬ (oscillating) and extract the sub-blocks,

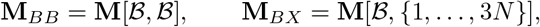

and then store (*µ***M**_*BB*_)^−1^. We note that the pre-assembly accelerates the computation significantly. The target boundary velocity is

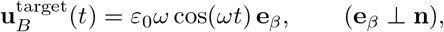

where **n** is the plate normal. For each evaluation of the right hand side, we first assemble the internal spring forces **g**_int_ and then solve for the unknown boundary tractions

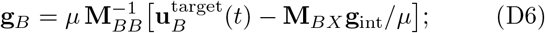

Finally, we obtain all node velocities **u** = *µ*^−1^**M** (**g**_int_ + **g**_*B*_). Spring rest lengths are updated through Eq. (D3) and the state is also advanced with dopri5. The time step Δ*t* is chosen such that *ω*Δ*t* ≤ 1*/*500; two periods are simulated and the first is discarded. We note that for large *E* and *η* values at small frequencies, the solver can be slow and unstable. Therefore, we set Δ*t* = 0.01 to obtain low-frequency results in Fig. 11b and e.

The instantaneous shear stress on the driven plate is *σ*(*t*) = (∑_*i*∈ℬ_ **g**_*i*_) · **e**_*β*_*/A*. Storage and loss moduli are extracted using the trapezoidal rule from the last period *T* = 2*π/ω* by [84]

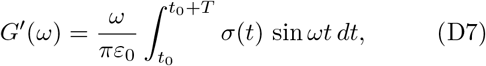

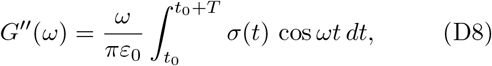

## Appendix E: Response of a single Jeffreys Element to creep and relaxation tests

We consider a single Jeffreys element described by the constitutive relation in Eq. 2. A step stress pulse of amplitude *σ*_0_, which turns on at time *t*_0_ and off at *t*_1_:

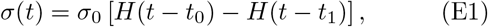

where *H*(*t*) denotes the Heaviside step function.

To solve for the strain *ε*(*t*), we take the Laplace transform of both sides of Eq. (2). We assume the system is initially at rest:

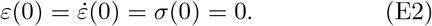

Substituting into Eq. (2):

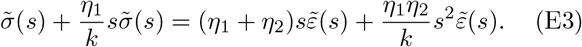

Rewrite with *λ*_1_ = *η*_1_*/k, λ*_2_ = *η*_1_ + *η*_2_ and *λ*_3_ = *η*_1_*η*_2_*/k* and solve for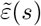:

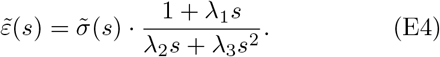

Substitute the Laplace transform of the stress pulse from Eq. (E1):

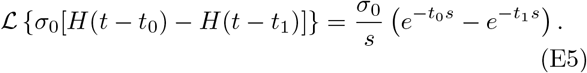

Substitute Eq. (E5) into Eq. (E4):

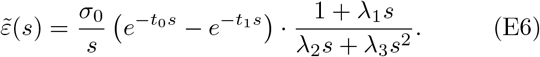

Finally, inverse Laplace transform to obtain the time-domain solution:

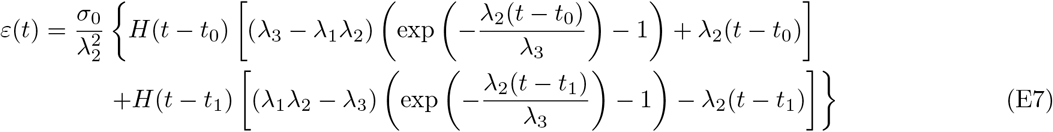

Next, for a stress relaxation test, we apply a step strain [84]:

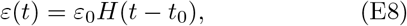

and we define the relaxation modulus as:

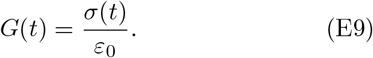

Similarly, taking the Laplace transform and solving analytically, we obtain:

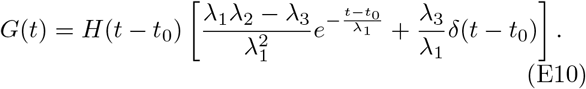

## Appendix F: Effect of uniform dilation on a Jeffreys network

Following the definitions in Appendix C, we consider a discrete viscoelastic network of *N* nodes at positions {**x**_*i*_} and edges ℰ= {(*i, j*)}with current lengths 𝓁_*ij*_ = ∥**x**_*i*_− **x**_*j*_∥. Each edge behaves as a one-dimensional Jeffreys element whose force-extension law in the frequency domain is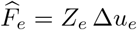, where *Z*_*e*_ follows Eq. C1.

Apply a purely geometric scale factor *s >* 0:

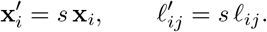

Because all coefficients in Eq. C1 scale as 1*/*𝓁_*e*_,

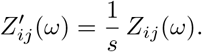

In the small amplitude oscillatory shear test used throughout this work, we drive the top slab by a sinusoidal displacement 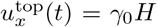 sin *ωt* on the original network, keeping the bottom slab fixed. After dilation the gap becomes *H*′ = *sH* but the same displacement amplitude 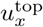 is applied. Hence

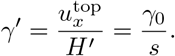

Node displacements are unchanged, so every edge experiences the same complex extension: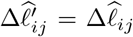. One has

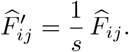

Summing over all edges cut by the top plane gives the total reaction force 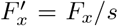. The top area dilates as *A*′ = *s*^2^*A*, so the shear stress becomes

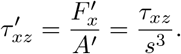

With *γ*_0_ unchanged, the complex shear modulus scales as

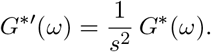

It shows that a uniform dilation softens both storage and loss moduli by *s*^−2^, reflecting the reduced edge density per unit area. To keep moduli size-independent one must refine the lattice (increase the number of edges per unit area) as the specimen is enlarged, ensuring a well-defined continuum limit.

## Appendix G: Random Geometric Graph inside a spherical domain

Random geometric graphs have been discussed by Dall and Christensen [135]. The properties of random graphs on the spherical surface were examined by Allen-Perkins [136]. Here we consider RGGs inside a spherical domain in ℝ^*d*^.

Consider a set of *n* nodes that are uniformly distributed in a domain *D* with volume *V* (*D*). Two nodes are connected by an edge if the Euclidean distance between them is less than or equal to a cutoff distance *r*. We define the indicator random variable *I*_*ij*_ for each unordered pair (*i, j*) as

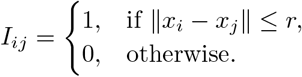

Then the total number of edges *M* in the graph is

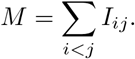

Taking the expectation, by linearity, we obtain

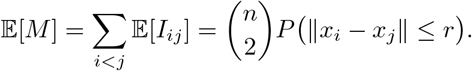

Since the nodes are uniformly distributed, the probability that two randomly selected points *x* and *y* are within a distance *r* is given by

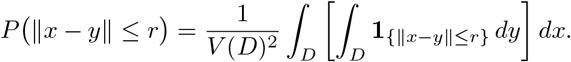

For sufficiently small *r* (compared to the size of the domain), the ball *B*(*x, r*) centered at *x* will be entirely contained in *D* for almost all positions of *x*. Therefore, the inner integral approximates to *V*_*d*_(*r*), and we get

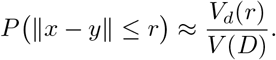

For a 3-dimensional spherical domain with radius *R*, substituting the expression for *P* (∥*x*− *y*∥≤ *r*) back into the expected number of edges, we have

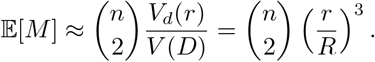

This can be generalized to higher dimensions. The volume of a *d*-dimensional ball of radius *r* in ℝ^*d*^ is given by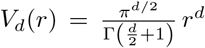, where Γ(·) is the Gamma function. Thus, in a general *d*-dimensional space the expected number of edges is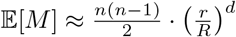.

## Appendix H: Monte Carlo Simulations using LaSSI

We set up simulations following the protocols described in Farag et al., [70]. Simulations with 200 molecules were identical to that of Farag et al., whereas simulations 2,000 macromolecules were performed anew. LaSSI is a lattice-based Monte Carlo simulation engine that is based on the generalized bond fluctuation model. The model for A1-LCD uses one lattice bead per amino acid residue. LaSSI can use either a transferable interaction potential, which on a lattice is a contact potential, or it can use bespoke potentials that are either phenomenological or learned for specific systems or sets of systems. Here, we use the bespoke potentials developed by Farag et al., [70, 71]. The binodal, which was calculated using temperature-dependent radial density profiles, were found to match the measured binodals for a range of systems including A1-LCD [70, 71]. The thermostat is exact, since LaSSI uses a Monte Carlo simulation engine, where moves are proposed to ensure preservation of detailed and accepted / rejected based on the Metropolis-Hastings criterion. Total system energies were calculated using a nearest neighbor model whereby any two beads that are within one lattice unit of each other along all three coordinate axes contribute to the total energy of the system. The full set of Monte Carlo are those of Farag et al., [70].

To extract the network from dense phases, we used radial density profiles to delineate the coexisting dense and dilute phases. The coordinates of chain residues on a lattice were converted into graph-theoretic descriptions of networks using the nearest neighbor adjacency on a the lattice. This checks each of the aromatic residues on each A1-LCD chain for contacts with the nearest 26 neighboring voxels. If a pairt of aromatic residues from two different chains occupy an adjacent voxel, then the pair of chains are considered to be neighbors with an edge between them.

To convert from volume fraction to mass concentration, which enables comparison between other simulation methods, we followed Wei et al., [49] and Farag et al., [70]. A volume fraction of 1.0 corresponds to a mass concentration of 1,310 mg/ml. Following Farag et al., [70], we used a multiplicative factor of 5.6 to convert the simulation temperature in LaSSI (which is written in units of *k*_*B*_ = 1 to degree Kelvin. This factor was derived as follows: Binodals extracted from LaSSI simulations were converted into mass concentrations along the abscissa and the ordinate was multiplied by a conversion factor that maximized the overlap with the measured binodals for A1-LCD. This conversion, which was derived for A1LCD, was then applied without further modification, to more than 30 different sequences derived from the A1LCD parent [70] and to the LCD of FUS[71]. Here, we used the factor of 5.6 as prescribed by Farag et al. The conversions from volume fractions to mass concentrations and simulation temperatures to Kelvin allow us to set up equivalent and comparative off-lattice molecular dynamics simulations using Mpipi-Recharged and CALVADOS.

## Appendix I: Mpipi-Recharged simulations

We used the Mpipi-Recharged built on OpenMM [137] called OpenMpipi [76]. To ensure equivalency with the LaSSI simulations for 200 macromolecules in a cubic box with side length 120 lattice units, the effective concentration was estimated to be ~ 20.77 mg/ml. Instead of the conventional approach of using slabs for simulation cells, we performed simulations (Fig. 7) for 200 A1-LCD molecules using cubic boxes with the target density of 20.77 mg/ml with periodic boundaries. This yielded a simulation box with the length of each side being 593.6 Å. The simulations were initiated by applying a compressive restraining force that draws all A1-LCD molecules to the center of box. This was followed by energy minimization to remove steric overlaps and quench the system into a single, densely packed droplet. The restraint was then removed, and the system was relaxed via thermal molecular dynamics simulations to achieve two-phase equilibrium. Equilibration was monitored by the convergence of the potential energy of the overall system and by equilibration of the density of the dense phase. To enable the simulations, as designed, and to extract network information from OpenMpipi, certain modifications were needed to the original code base. These modifications can be accessed in the forked repository (https://github.com/zhangruoyao68/OpenMpipi).

To extract graphs from snapshots, we adopted the following protocol: The boundary between dense and dilute phases was first delineated. Next, the average inter-residue distance (*r*_res_ ≃ 3.80 Å) for all chains was evaluated. The cutoff distance for identifying edges was set to 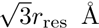 to match the cutoff factor used in the LaSSI networks. Edges were drawn from chains that had residues that matched the criterion of inter-residue contacts between chain molecules. Note that our goal was not to calculate converged binodals. Instead, our goal was to extract snapshots of dense phases that are representative of the converged ensemble of configurations sampled in Mpipi-Recharged for the A1-LCD system.

## Appendix J: CALVADOS simulations

We also used CALVADOS to produce droplet configurations [138]. The overall approach, with a few CALVADOS-specific modifications, follow the approach we used for Mpipi-Recharged. We set the box size to be 593.6 Å to match the mass concentrations in the Mpipi-Recharged and LaSSI simulations. The protocol was otherwise the same as that used for Mpipi-Recharged. Slight modifications of the output format were needed to extract network information from CALVADOS, and these can be found in the forked repository (https://github.com/zhangruoyao68/CALVADOS). Graphs were derived using contact criteria that were identical to what was used for Mpipi-Recharged.

## Appendix K: Visualization

The output files of network structures and edges in .vtk/.vts format were written using python package PyVista [139]. Visualization of snapshots and graphs were performed using ParaView [140]. The structures of the coarse-grained biomolecular condensates were rendered using OVITO [141].

